# Quantitative modeling of pentose phosphate pathway response to oxidative stress reveals a cooperative regulatory strategy

**DOI:** 10.1101/2022.02.04.478659

**Authors:** Julien Hurbain, Quentin Thommen, Francois Anquez, Benjamin Pfeuty

## Abstract

Living cells use signaling and regulatory mechanisms to adapt to environmental stresses. In the case of oxidative stress due for instance to hydrogen peroxide exposure, the adaptation response relies on co-regulation of enzymes in both glycolysis and pentose phosphate pathways (PPP), so as to support PPP-dependent *NADPH* and redox homeostasis. To understand the regulatory logic underlying early oxidative stress response, available metabolomics and ^13^C fluxomics dataset are used to infer a probabilistic ensemble of kinetic models. Model ensemble properties of parameter distributions, transient dynamics, dose-response curves and loss-of-function phenotypes all highlights significant and cooperative effects of allosteric regulations of G6PD, PGI and GAPD in early oxidative response. Indeed, efficient flux rerouting into PPP is shown to require dose-dependent coordination between upregulated G6PD enzyme and increased G6P metabolite, the latter requiring fine-tuned inhibition of upper and lower glycolytic enzymes. This set of allosteric regulation also combines negative and positive feedback loops in a subtle manner prone to generate paradoxical perturbation phenotypes for instance related to 6PGD modulation.

## Introduction

The oxidative pentose phosphate pathway (OxPPP) is a fundamental pathway of glucose metabolism involved in nucleotid biosynthesis and redox homeostasis (Stincone *et al*., 2015). Its role is prominent following an oxidative stress to generate *NADPH* required for fueling anti-oxidant machinery and producing biosynthetic precursors required for repairing DNA damages. A significant increase of metabolic flux in this pathway is commonly observed in any living cells subjected to oxidative stress (Ben-Yoseph *et al*., 1996; Ralser *et al*., 2007; LaMonte *et al*., 2013; Kuehne *et al*., 2015; Christodoulou *et al*., 2018; Nikel *et al*., 2021). A common rationale for such metabolic flux rerouting relies on the acknowledged roles of *NADP* as a coenzyme and *NADPH* as a competitive inhibitors of the first oxidation reaction of the OxPPP (Warburg and Christian, 1936; Negelein and Haas, 1935; Eggleston and Krebs, 1974). The scavenging activity of glutathione antioxidant is coupled to the oxidation of *NADPH* into *NADP* ^+^, which is therefore prone to increase *G*6*PD* activity and *NADPH* production. However, oxidative stress has also been shown to induce directly or indirectly the allosteric inhibition of diverse glycolytic enzymes such as *PGI* (Kuehne *et al*., 2015; Dubreuil *et al*., 2020), *GAPD* (Ralser *et al*., 2007; Peralta *et al*., 2015), *PK* (Anastasiou *et al*., 2011) or *TPI* (Grüning *et al*., 2014). A complex pattern of regulation at the levels of PPP and glycolysis raises the question of their coordination for efficient metabolic rerouting.

The metabolic network combining upper glycolytic and pentose phosphate pathway displays a complicate branching structure comprising both reversible and irreversible reactions, which obstructs intuitive understanding of multisite metabolic regulation. To investigate complex regulatory patterns, the kinetic modeling framework is often used to disentangle the respective and cooperative roles of multiple feedback regulations (Muzzey *et al*., 2009; Relógio *et al*., 2011; Pfeuty *et al*., 2018; Sander *et al*., 2019). Several kinetic models have addressed OxPPP dynamics with respect to a specific organism and experimental dataset (Thorburn and Kuchel, 1985; Schuster and Holzhütter, 1995; Kerkhoven *et al*., 2013). Nowadays, advanced metabolomics and fluxomics studies such as kinetic measurements of concentrations and isotopic labeling patterns provide a rich material to build increasingly reliable and comprehensive kinetic models (Miskovic *et al*., 2015; Foster *et al*., 2019; Hameri *et al*., 2019). Regarding the regulation of the oxidative stress response, such data are available and have already been analyzed in terms of significance and ranking of diverse regulatory hypothesis (Kuehne *et al*., 2015, 2017; Christodoulou *et al*., 2019). However, questions remain about the interplay and cooperativity between those different regulatory mechanisms.

In the present study, we aim to build a class of kinetic models inferred from a comprehensive dataset associated with the oxidative stress response of fibroblast (Kuehne *et al*., 2015). On the one hand, the network structure of the model is chosen to fit with the available data and with the objective to understand metabolic flux rerouting for *H*_2_*O*_2_ detoxification. On the other hand, flux analysis from ^13^C-labeling data and parameter estimation from flux and concentration data are based on Monte Carlo sampling methods (Saa and Nielsen, 2016; St. John *et al*., 2019; Valderrama-Bahamóndez and Fröhlich, 2019; Theorell and Nöh, 2020) so as generate a representative sample of kinetic models consistent with experimental measurement values and their uncertainties. From such model ensemble, we perform a comprehensive set of analysis regarding parameter distributions, transient dynamical responses, doseresponse properties and gain/loss-of-function phenotypes, which portrays the manner how allosteric regulations contribute to the metabolic response upon oxidative stress. In particular, these analysis converge to the notion that distributed allosteric regulation is required for efficient metabolic rerouting where regulatory mechanisms display both complementary and cooperative roles.

## Result

### Kinetic modelling approach for metabolic response to oxidative stress

We design a metabolic model whose specificities match with the particular context of the early metabolic response to oxidative stress (Figure 1A). So far, kinetic models of oxidative stress response investigate early intra-cellular responses either at the level of antioxidant pathways (Adimora *et al*., 2010; Kembro *et al*., 2013; Benfeitas *et al*., 2014) or at the level of metabolic glycolysis and PPP pathways (Thorburn and Kuchel, 1985; Schuster and Holzhütter, 1995; Kerkhoven *et al*., 2013; Christodoulou *et al*., 2018). The latter studies were either considering passive metabolic response without regulation or exploring all possible metabolic-enzyme interactions. In contrast, our model specifically addresses the respective role of a subset of regulatory mechanisms that has been acknowledged to contribute to some respect to rapid metabolic adaptation to oxidative stress (Mullarky and Cantley, 2015; Dick and Ralser, 2015), namely *NADPH*-dependent inhibition of PPP enzymes (Yoshida and Lin, 1973; Christodoulou *et al*., 2018), 6*PG*-dependent inhibition of *PGI* (Kahana *et al*., 1960; Gaitonde *et al*., 1989; Kuehne *et al*., 2015; Dubreuil *et al*., 2020), *H*_2_*O*_2_-dependent inhibition of *GAPD* (Ralser *et al*., 2007; Peralta *et al*., 2015; van der Reest *et al*., 2018) and regulation of *NADPH*-consuming or -producing reactions (Fan *et al*., 2014; Jiang *et al*., 2016; Chen *et al*., 2019). The dynamics of the metabolic network including the selected pathways and regulations shown in Figure 1A is described by the differential equation system:

**Figure 1:**
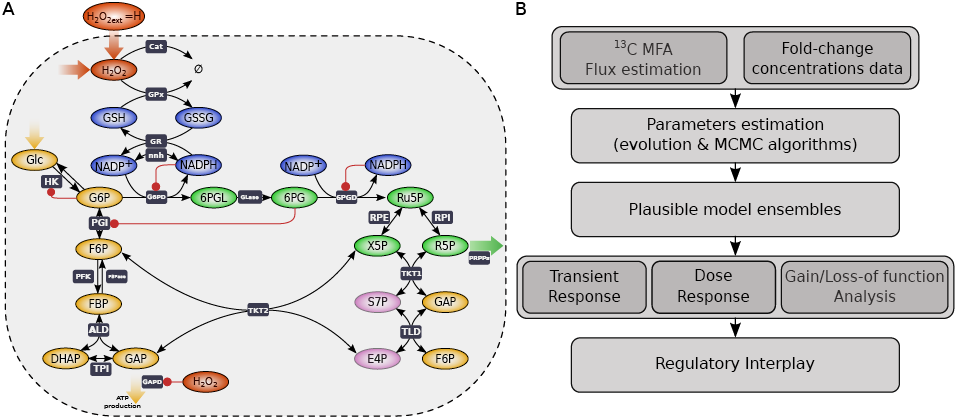
Modeling workflow. (A) Metabolic network module considered to study early metabolic response to oxidative stress. The network comprises the gluthatione, glycolytic and pentose phophate pathways, supplemented with a selected set of allosteric regulations (red arrows). (Glucose *Glc*; Glucose-6-phosphate *G*6*P* ; Fructose-6-phosphate *F* 6*P* ; Fructose-1,6-bisphosphate *FBP* ; Dihydroxyacetone phosphate *DHAP* ; Glyceraldehyde-3-phosphate *GAP*, 6-Phosphogluconolactone 6*PGL*; 6-phosphogluconate 6*PG*; Ribulose 5-phosphate *Ru*5*P* ; Xylulose-5-phosphate *X*5*P* and Ribose-5-phosphate *R*5*P* ; Sedoheptulose-7-phosphate *S*7*P* ; Erythrose-4-phosphate *E*4*P*). (B) Modeling workflow from 13C fluxomics and metabolomics data (Kuehne *et al*., 2015) to kinetic model ensemble, ending with regulation analysis of transient response, dose response and gain/loss-of-function phenotypes

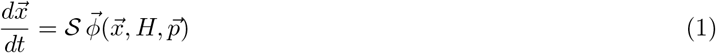

where 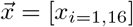 represents the concentrations of metabolite species 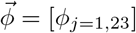 represents the reaction rates associated to enzymes *j*, 𝒮 is the stoichiometry matrix, *H* is the extracellular *H*_2_*O*_2_ and 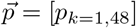 represents the enzymatic or regulatory parameters (see Supplementary Material for details). The steady-state equation is given by 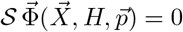 where capitalized 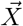 and 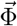 denote steady-state concentration and flux vectors.

Within a kinetic modeling framework, the strategic workflow to study the metabolic flux rerouting response following oxidative stress will proceed in several steps (Figure 1B): (i) Collecting and processing 13C labeling and metabolomics data (Kuehne *et al*., 2015), (ii) model parameter estimation of a kinetic model (using genetic and Monte-Carlo algorithms) to generate a plausible model ensemble and (iii) comprehensive analysis of these model ensembles to decipher the regulatory interplay for stress-induced metabolic flux rerouting.

### Refined analysis of metabolic flux rerouting

The stress-induced redistribution pattern of metabolic fluxes can be inferred without knowledge about kinetic parameters. Although an expected feature of such redistribution is the increased flux in the oxidative branch of PPP, it remains unclear to which extent is such increase and whether the OxPPP flux is rather directed toward nucleotide production or toward the nonoxidative branch of the PPP. To gain a quantitative description of oxidative stress-induced redistribution of metabolic fluxes, we reanalyse ^13^C labelling data (Kuehne *et al*., 2015) using SSA-^13^CMFA to simulate the isotope labeling system (Thommen *et al*., 2022) and Monte Carlo (MC) sampling to determine posterior distribution of flux parameters. We obtain a distribution of flux for all reactions (Figure 2A and S1A) associated with an accurate fit of mass itopomer data (Figure S1C,D). The size of confidence intervals associated to flux distribution (Figure S1B) determines whether the flux can be estimated accurately enough to be used as a constraint for kinetic modeling.

**Figure 2:**
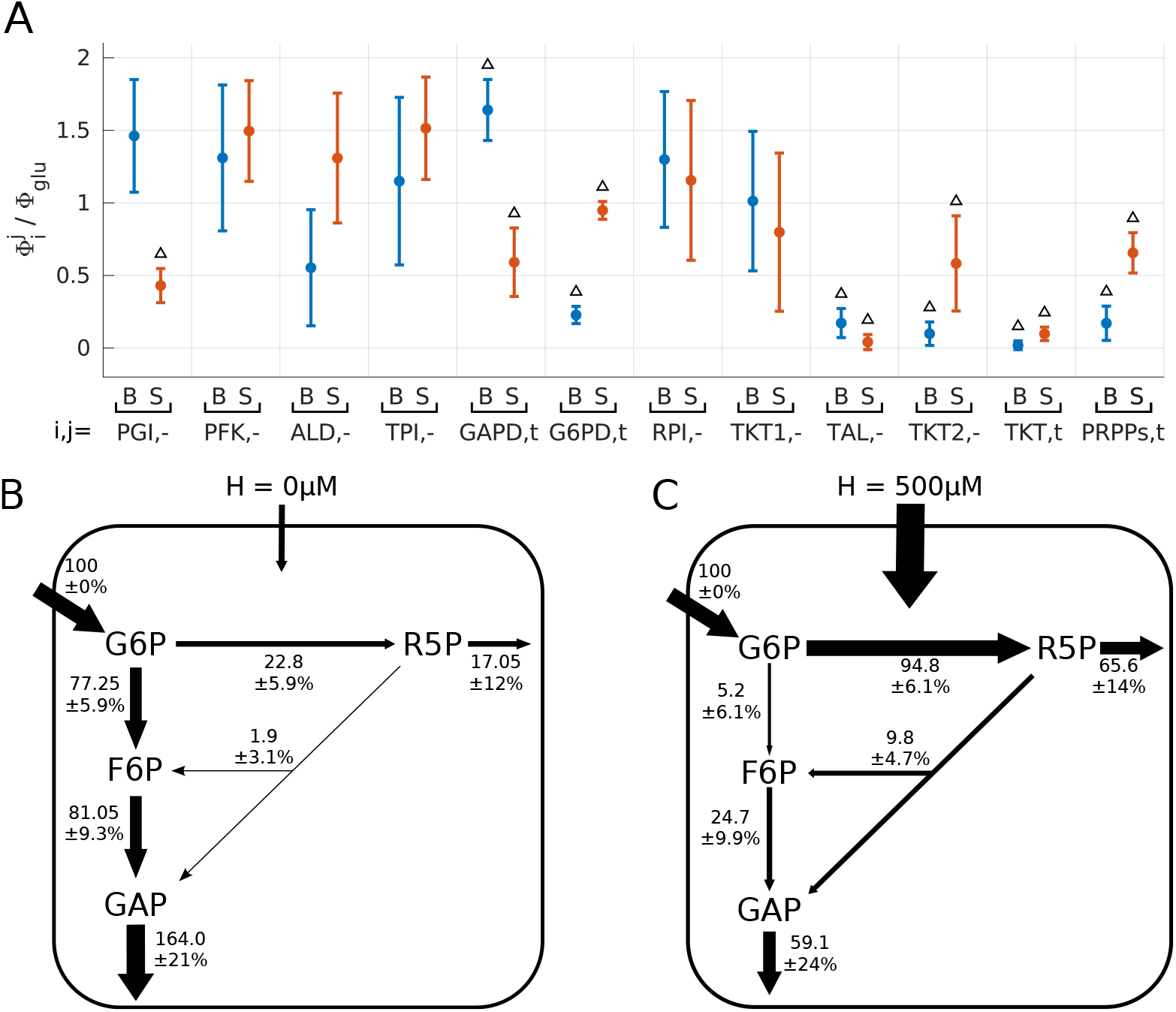
Stress-induced flux redistribution. (A) Mean and standard deviation of the distribution of normalized flux parameters obtained using SSA-based ^13^C-MFA. Estimation are restricted to a set of elementary flux parameters (while other fluxes can be derived from balance equations) where the index *i* indicates the enzyme and the index *j* indicates the directionality (+ are directed toward GAP), and where estimation is performed for basal (B, blue) or stress (S, red) conditions. Triangles indicate parameter estimation that are statistically-significant based on the relative size of confidence interval distribution (Figure S1B). (B-C) Estimation of the flux state in basal (B) and stress (C) conditions.

The estimated flux distribution pattern in absence and presence of oxidative stress can be summarized for the main branches of the metabolic network (Figure 2B-C). The metabolic state in the unstressed condition corresponds to a glycolytic flux mode where a minor fraction (∼ 20%) of glucose import flux is diverted toward OxPPP (Figure 2B). From such basal state, exposure of 500*µ*M of extracellular hydrogen peroxide leads to a significant increase of OxPPP flux to ∼ 95% which is further splitted toward nucleotid production and nonoxidative PPP (Figure 2C). In addition to the net fluxes associated to the branching architecture of the metabolic network, it is to note that some directional flux could be estimated in the nonoxidative PPP reactions as well as in the reversible *PGI* reaction, which are valuable informations for kinetic model building.

### Optimization and inference methods identify a plausible ensemble of kinetic models

Besides the redistribution of metabolic fluxes, oxidative stress response also induces rapid changes in metabolic concentrations at minute timescale (Kuehne *et al*., 2015), which together provides a valuable dataset to estimate the parameters of kinetic models described by Equation 1. Our parameter estimation problem consists in estimating the values of 36 parameters (as 12 parameters are fixed including equilibrium constant) from a dataset of the early metabolic response comprising 13 estimated values of fluxes in basal and stress conditions, and 12 measured values of concentration ratio. The procedure combines two classes of global optimization methods, namely an evolutionary genetic algorithm and a MCMC algorithm (see Materials and Methods), following a stepwise strategy recapitulated in Figure 3A. First, the *nRMSE* (Eq 3) of models whose parameters are randomly sampled (10^6^ runs) lies with a 50% confidence interval between 55.4 and 2.6, thus confirming the need of a dedicated parameter optimization procedure to find parameter model consistent with experimental data. Second, an evolutionary genetic algorithm is used as a preliminary step to generate a sample of optimized models whose 50% confidence interval *nRMSE* values lies between 3 and 0.8. Third, a small subset of such local optimum solutions whose *nRMSE*(*p*) < 1 is used as initial conditions of MCMC sampling algorithm of the parameter space. The parameter distribution obtained with large enough sampling of ∼ 10^6^ accepted steps (i.e., smooth stationary distribution for most parameters as shown in Figure S2A) provides an estimation for parameter uncertainty and defines a statistical ensemble of kinetic model that is called 𝒫^*opt*^ and that is analyzed in details in the following. Such statistical model ensemble is characterized with a 50% confidence interval of *nRMSE* between 0.63 and 0.86 for which the estimated distributions of flux and concentration ratio fall within the range of experimental uncertainties (Figure 3B-C). Parameter distributions shown in Figure 3D discriminate between stiff and sloppy parameters for which 50% confidence intervals span from few percents to several order of magnitude of parameter variations. Spectral analysis of correlation matrix confirms indeed the existence of a few poorly estimated parameters generally associated with strong correlation between parameters of a same reaction (Figure S2B,C). Last but not least, a dataset about dynamic and dose-dependent concentration responses is available (Kuehne *et al*., 2015) and has been retained to assess the predictive capability of the plausible set of model 𝒫^*opt*^ (Figure S3). The timescale and threshold of concentration response are overall well predicted, except sparse discrepancies (e.g., lower threshold of 6*PG* response) that can be solved by adding these data in the score function.

**Figure 3:**
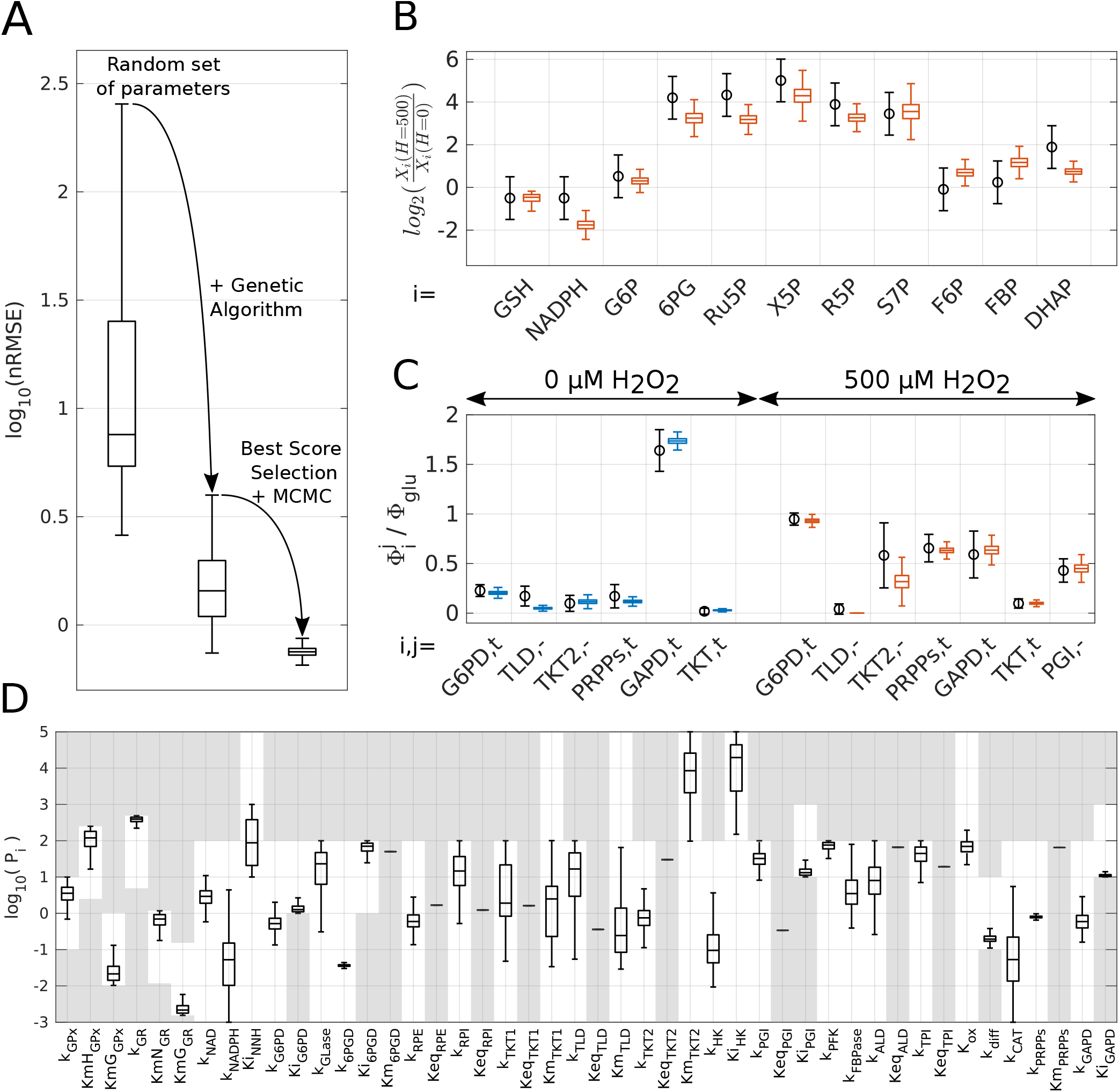
Model selection and parameter estimation. (A) Whisker plots associated to a random parameter set (10^6^), an optimized parameter set using evolutionary genetic algorithm (10^4^) and a parameter set (𝒫^*opt*^) sampled with MCMC algorithm (10^6^). (B,C) Whisker plots of the concentration ratio *X*_*i*_(*H* = 500)*/X*_*i*_(*H* = 0) for the model ensemble 𝒫^*opt*^ as compared with the mean and standard deviation of experimental values (blue). (C) Whisker plots of a subset of normalized reaction fluxes Φ_*i*_*/*Φ_*GLU*_ for the model ensemble 𝒫^*opt*^ as compared with the mean and standard deviation of estimated values (blue). (D) Whisker plots of parameters for the model ensemble 𝒫^*opt*^ highlighting that parameter exploration is bounded to the white (non-shaded) parameter space

In summary, the parameter estimation procedure generates a plausible set of kinetic models whose parameters show rather narrow distributions, except some parameters that have a little impact on data adjustement or that can be compensated by change of other parameters. Importantly, most regulation parameters (*Ki*_*G*6*P D*_, *Ki*_*GAP D*_ and *Ki*_*P GI*_) shows a narrow distribution, confirming the role of these regulations in shaping the metabolic response to oxidative stress. In the following, we systematically perform analysis on this model ensemble 𝒫^*opt*^ to draw a statistical picture of the regulatory properties.

### Transient dynamics during stress response displays a multiphasic time course

The characteristics of the transient dynamics induced by the oxidative stress before reaching steady state gives some preliminary insights about the respective contribution of passive and regulated metabolic response (Figure 4). The minute time resolution in the time series dataset seemed not sufficient to identify trends arising at second timescale (Christodoulou *et al*., 2018). In simulation of the model ensemble 𝒫^*opt*^, metabolite species within a same metabolic module share a similar dynamic response profile (Figure 4). First, PPP metabolites display a rapid and significant monophasic increase. Second, detoxifying *NADPH* and *GSH* metabolites display a fast and significant decrease followed by a slower increase. Third, Glycolytic metabolites show moderate changes where a fast decrease seems followed by a slower increase. These different temporal response patterns are typically characterized with a biphasic response where a fast passive response to perturbation is quickly followed by a slower regulated response. The biphasic nature of the transient response is illustrated in the case intracellular concentration of *H*_2_*O*_2_ and *G*6*P* (Figure 4B,C). *H*_2_*O*_2_ shows a sharp increase about several order of magnitude whose timescale within seconds that relates to the basal degradation timescale *k*_*GR*_ × *KmG*_*GP x*_ ∼ 10s. The time profile of *H*_2_*O*_2_ later displays an overshoot in the time course where decrease of *H*_2_*O*_2_ mirrors the increase of *GSH* and *NADPH*, which coincides with the upregulation of glycolytic metabolites including *G*6*P*. *G*6*P* shows indeed a rapid decrease until ∼ 10s before to increase again to eventually exceeds its initial value (Figure 4C).

**Figure 4:**
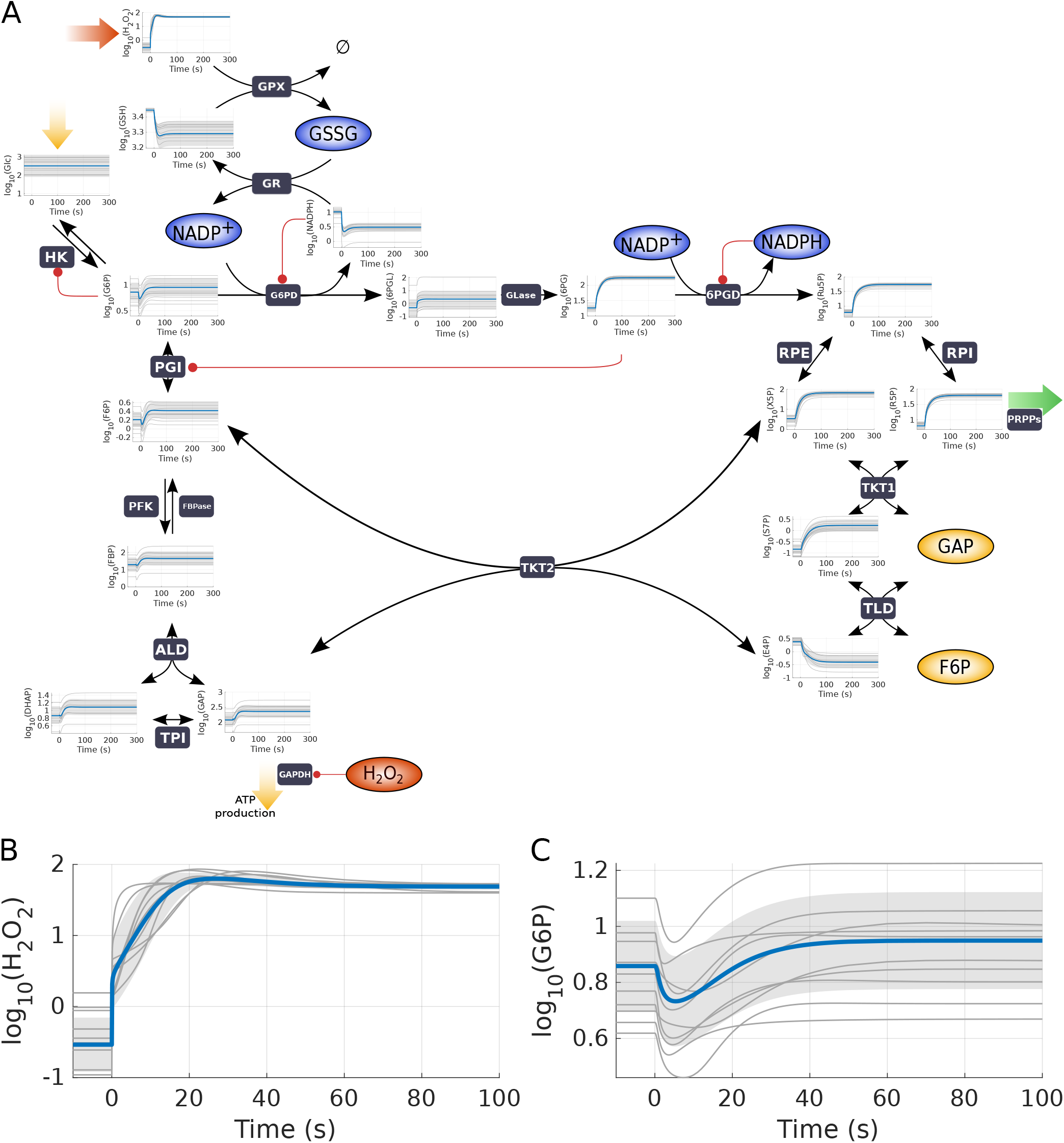
Portrait of transient dynamic responses. Statistical picture of the transient dynamic response of metabolites concentrations to in response to a step of 500*µ*M *H*_2_*O*_2_. Plots of mean value (blue line), a subsample of 50 trajectories (grey line) and the standard deviation (grey shadow) for a subsample of 10^5^ parameter set sample 𝒫^*opt*^ (10^5^ parameter set) (A) Dynamic response of all metabolite species until 5mn. (B,C) Dynamic response of *H*_2_*O*_2_ and *G*6*P* until 100s, highlighting the multiphasic time course.

The early decreasing phase of *G*6*P* dynamics coincides with an increased consumption through *G*6*PD* associated to higher *NADP* ^+^ levels while the late increasing phase can only be due to a decreased glycolytic flux through *PGI* probably related to allosteric inhibition of glycolysis at the level of *GAPD* or *PGI*.

In summary, the biphasic response observed in simulations of the plausible set of models distinguishes between a detoxification response in less than seconds using the reservoir of glutathione and cofactor *NADPH* and a metabolic rerouting response in tenth of seconds that involves inhibition of glycolysis to quickly restore high G6P levels.

### Dose-response analysis identifies rate-limiting reactions

The redistribution pattern of metabolic fluxes is determined for a specific level of *H*_2_*O*_2_ for which ^13^C labeling data were available. The OxPPP flux normalized to glucose import flux, Φ_*G*6*P D*_*/*Φ_*GLU*_, is distributed around 1 (Figure 2A) which is far below its maximal flux capacity associated to full inhibition of *GAPD* and *PRPP* enzyme (i.e., Φ_*G*6*P D*_*/*Φ_*GLU*_ ≤ 6). We therefore perform a dose-response analysis of the kinetic model ensemble 𝒫^*opt*^ focusing respectively on the antioxidant response, concentration response and flux response (Figure 5). Simulation of the metabolic response at 5mn as function of the extracellular level of *H*_2_*O*_2_ shows a transition in the detoxification response around 500*µM* (Figure 5A). Below this value, the detoxification activity of *GPx* increases with oxidative stress level thereby keeping low intracellular levels of *H*_2_*O*_2_ at the expense of an increased reduced state of glutathione. Above this value, *GPx* activity saturates such that metabolic flux through this reaction is bounded and cannot increase anymore to compensate for the increase of *H*_2_*O*_2_ production above some level. Beyond this threshold, *H*_2_*O*_2_ is eliminated through catalase, consistently with the idea of that the rate-limiting enzymes depend on intracellular *H*_2_*O*_2_ concentrations (Makino *et al*., 2004; Ng *et al*., 2007). Another qualitative change of the metabolic response is observed for large enough *H*_2_*O*_2_ (Figure 5B). 6*PG* does not reach a steady state and continue to slowly increase, as values differ between 5mn and 30mn. The appearance of a slower equilibration dynamics of 6*PG* coincides with the saturated kinetics of 6*PGD* enzymatic reaction associated to increased levels of 6*PG* below that value of *Km*_6*P GD*_. In parallel to the concentration changes of *H*_2_*O*_2_ and 6*PG*, the dose response of metabolic fluxes displays a gradual change from a glycolytic mode to OxPPP mode up to a rather cycling mode (Φ_*G*6*P D*_*/*Φ_*GLU*_ ∼ 1.3 > 1) (Figure 5C).

**Figure 5:**
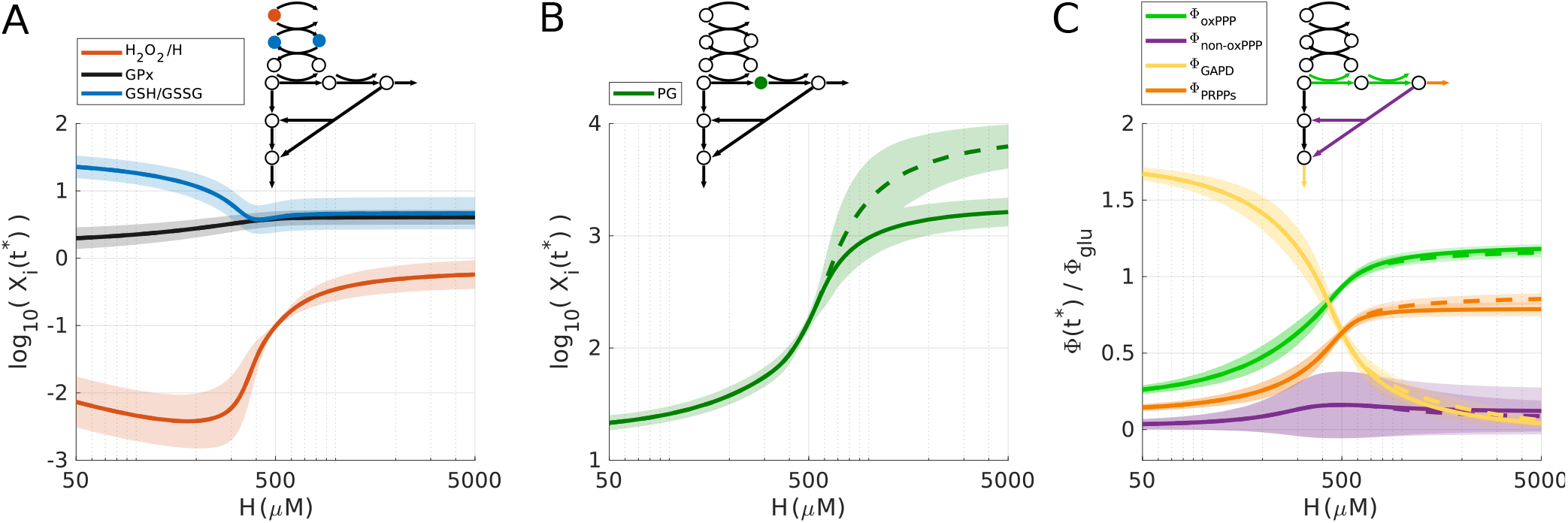
Dose-dependent profile of the oxidative stress response. Metabolic response to varying level of extracellular *H*_2_*O*_2_ measured at 5mn (full line) and 30mn (dashed line). (A) Dose response in the gluthatione pathway of 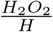, *GPx* and *GSH*/*GSSG* ratio. (B) Dose response of 6*PG* metabolite (other metabolites are shown in Figure S4). (C) Dose response of the elementary fluxes through enzymatic reactions *G*6*PD, TKT, GAPD* and *PRPP*, and normalized to glucose import rate *ϕ*_*GLU*_.

In conclusion, dose-response analysis identifies several rate-limiting mechanisms, where saturation in *GPx* restrains *H*_2_*O*_2_ detoxification, while saturation in 6*PGD* and leak flux through PRPPs reactions restrains the maximal flux into OxPPP for NADPH homeostasis. Naively, this limited flux recycling can be interpreted as a need for nucleotid production for DNA repair and could be relieved through transcriptional regulation mechanism.

### Regulation analysis reveals both complementary and cooperative regulatory interplay

The model comprises several regulatory mechanisms that can contribute to the metabolic flux rerouting to oxidative stress. Some of these regulations directly act at the levels of reactions producing or consuming NADPH notably in OxPPP (*Ki*_*G*6*P D*_, *Ki*_6*P GD*_, *Ki*_*NNH*_) while others operate at the level of glycolysis (*Ki*_*GAP D*_, *Ki*_*P GI*_). The estimated values of *Ki* provide an initial insight for the need of regulation to match kinetic model with experimental data. Computation of inhibitory strength *X*_*j*_*/Ki*_*j*_ (= 1 for enzymatic activity divided by two) over the model ensemble 𝒫^*opt*^ reveals that stress condition is associated with significant inhibition of *PGI* and *GAPD* (*X*_*j*_*/Ki*_*j*_ ≫ 1) but also significant disinhibition of G6PD (Figure 6A).

**Figure 6:**
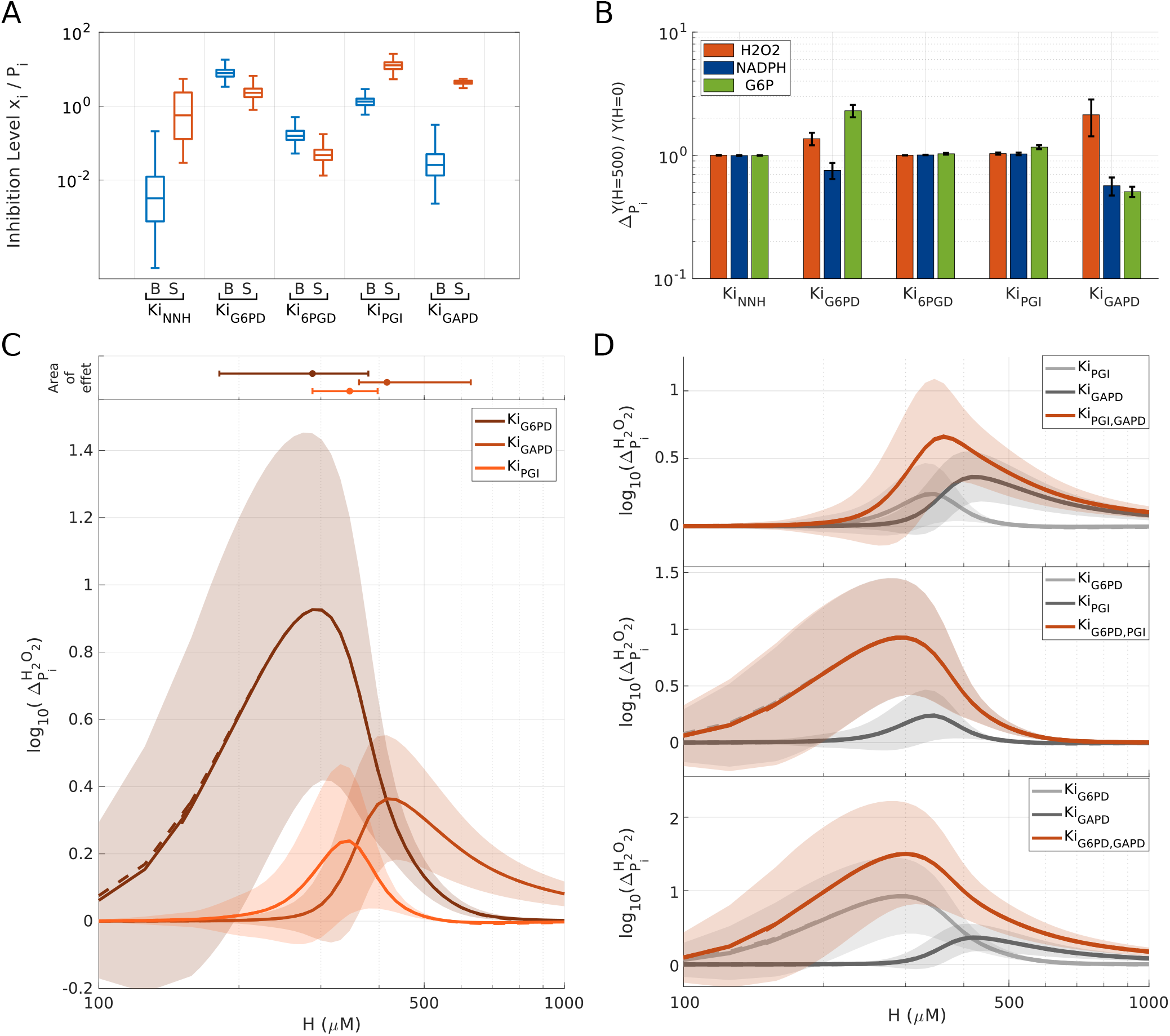
Regulation analysis based on gain/loss-of-function simulations. (A) Inhibitory strength *X*_*i*_*/Ki*_*i*_ of enzyme *i* associated to inhibition constant *Ki* in basal (blue) and 500*µM H*_2_*O*_2_ (red) conditions. *X*_*i*_*/Ki*_*i*_ = 1 indicates that inhibition reduces enzymatic activity by a factor 1 + *X*_*i*_*/Ki*_*i*_ = 2. (B) Sensitivity factor 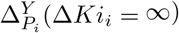 (Eq. 4) of output *Y* with respect to several deletions of regulation *Ki*_*i*_ in abscisse, where the outputs *Y* are the metabolic response of *H*_2_*O*_2_, *NADPH* and *G*6*P* to oxidative stress relative to basal levels. Bars are mean values and error bars are standard deviations over the kinetic model ensemble 𝒫^*opt*^. (C) Sensitivity factor 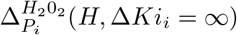 of *H*_2_*O*_2_ with respect to deleted regulation *Ki*_*GAP D*_, *Ki*_*G*6*P D*_, *Ki*_*P GI*_ as function of extracellular *H*_2_*O*_2_, *H*. Thedose-specific areas of regulatory effect associated to each deleted regulation (Δ(*H*)*/*Δ_*max*_ > 0.5) is shown upper to the panel. Sensitivity factor associated to key metabolite concentrations are shown in Figure S5. (D) Sensitivity factor 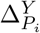 with respect to combined deletion of two regulations in comparison to the single deletions.

A more comprehensive strategy to quantify regulatory effects consists in measuring gain-of-function or loss-of-function associated to the deletion of one of those regulatory mechanisms (*Ki*_*i*_ → ∞), other things being equal. For such aim, we define a sensitivity quantity Δ measuring the impact of some parameter change (e.g., lower allosteric inhibition) on some functional output (e.g., intracellular *H*_2_*O*_2_) (Eq 4). 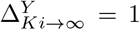 indicates that deleting a regulation *Ki* has no phenotypic effect while 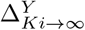 larger or lower to 1 indicates that deleting the regulation increases or decreases the steady-state level of *Y*. In Figure 6B, removing *NAPDH*-dependent inhibition of *G*6*PD* or *H*_2_*O*_2_-dependent inhibition of *GAPD* leads to higher *H*_2_*O*_2_ and lower *NADPH*, highlighting significant contribution of these regulations for NADPH homeostasis and *H*_2_*O*_2_ detoxification. More surprisingly, deleting other regulation whose strength is not necessarily negligible has a minor impact on metabolic outputs of interest. However, the absence of effect of deleting the 6PG-dependent inhibition of PGI can be interpreted by the reversibility of the enzymatic reaction where 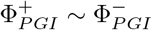 for this particular stress level (Figure 2C).

To investigate whether regulation efficiency depends on oxidative stress level, we evaluate how the deleterious effect of removing a regulatory link (i.e., loss-of-function) depends on extracellular hydrogen peroxide *H* (Figure 6C-D). The dose-dependent profile of 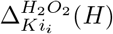 discriminates different ranges of stress level for which each regulation *Ki*_*i*_ is the most efficient (Figure 6C). First, the efficiency of G6PD upregulation is the highest for low-to-moderate stress level, probably because enzyme activity rather than G6P level is rate-limiting. Second, the efficiency of 6PG-dependent inhibition of PGI (null for *H* = 500 in Fig. 6B) peaks at intermediate stress level as it requires both (i) high enough stress for 6PG accumulation (see Fig. 5B) and (ii) not-too-high stress such that downstream flux prevails over upstream flux in PGI reaction (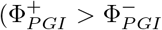 or equivalently Φ_*G*6*P D*_*/*Φ_*GLU*_ < 1) (see Fig. 5C).

Third, the efficiency of GAPD inhibition culminates at high oxidative stress consistently with its ability, in addition to restore G6P levels, to reverse glycolytic flux coming from the non-oxidative branch of the PPP, thus making possible a prevailing cycling mode where Φ_*G*6*P D*_*/*Φ_*GLU*_ > 1.

Besides complementary efficiency ranges, two regulations can also combine their effect in a non-trivial synergistic or redundant manner. For instance, upregulation of G6PD enzyme enhances 6PG metabolite levels which potentializes inhibition of PGI. Alternatively, inhibition of PGI activity reduces consumption of G6P so as to potentialize the upregulation of G6PD. These are second-order effects that are quantified by the computation of sensitivity factors 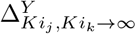 associated to the combined deletion of two regulations (Figure 6D).

*G*6*PD*, 6*PGD* and *TKT* enzymes are common targets of loss-of-function experiments to investigate oxidative stress response (Chan *et al*., 2013; Kuehne *et al*., 2015; Nóbrega-Pereira *et al*., 2016; Wan *et al*., 2017; Sun *et al*., 2019; Li *et al*., 2019; Dubreuil *et al*., 2020). We therefore perform simulations of our model ensemble 𝒫^*opt*^ while varying the enzymatic activity parameter from 10-fold reduction to 10-fold increase. We record both the mean of *H*_2_*O*_2_, the *NADPH*-producing OxPPP flux and some concentration metabolites for the model ensemble 𝒫^*opt*^ (Figure 7). Expectedly, increase (resp., decrease) of *k*_*G*6*P D*_ leads to a more (resp., less) efficient oxidative stress response related to subsequent change in the OxPPP flux of *NADPH* production (Figure 7A,B). In sharp contrast, modulation of *k*_6*P GD*_ leads to a more surprising and ambivalent metabolic response (Figure 7C,D). Depending on the level of stress and of modulation, we observe that (i) both increase or decrease of *k*_6*P GD*_ can weaken the oxidative stress response, and (ii) decrease of *k*_6*P GD*_ can both weaken or improve oxidative stress response. These ambivalent phenotypes relate to the dual effect of modulating *k*_6*P GD*_, on the G6PD enzymatic rate itself but also on the increase level of 6*PG* that feedback on PGI activity, G6P level and G6PD flux. Specifically, reducing *k*_6*P GD*_ can lead to an imbalance state associated to increased metabolic flux through 6PGD and decreased flux through G6PD, whose relative effect determines occurrence of loss or gain of function. Last, the modulation of *k*_*T KT*_ has a moderate effect on flux reprogramming restricted to higher stress level (Figure 7E,F), consistently with the dose-dependent increase of flux in nonoxidative PPP (Figure 5C) and its crucial role for enabling a cycling flux where *ϕ*_*G*6*P D*_*/ϕ*_*GLU*_ > 1. It is to note that some trends observed for the concentration response of glycolytic and PPP intermediates to modulation of *k*_*G*6*P D*_ and *k*_*T KT*_ (Figure S6) are consistent with knockdown experiments (Kuehne *et al*., 2015).

**Figure 7:**
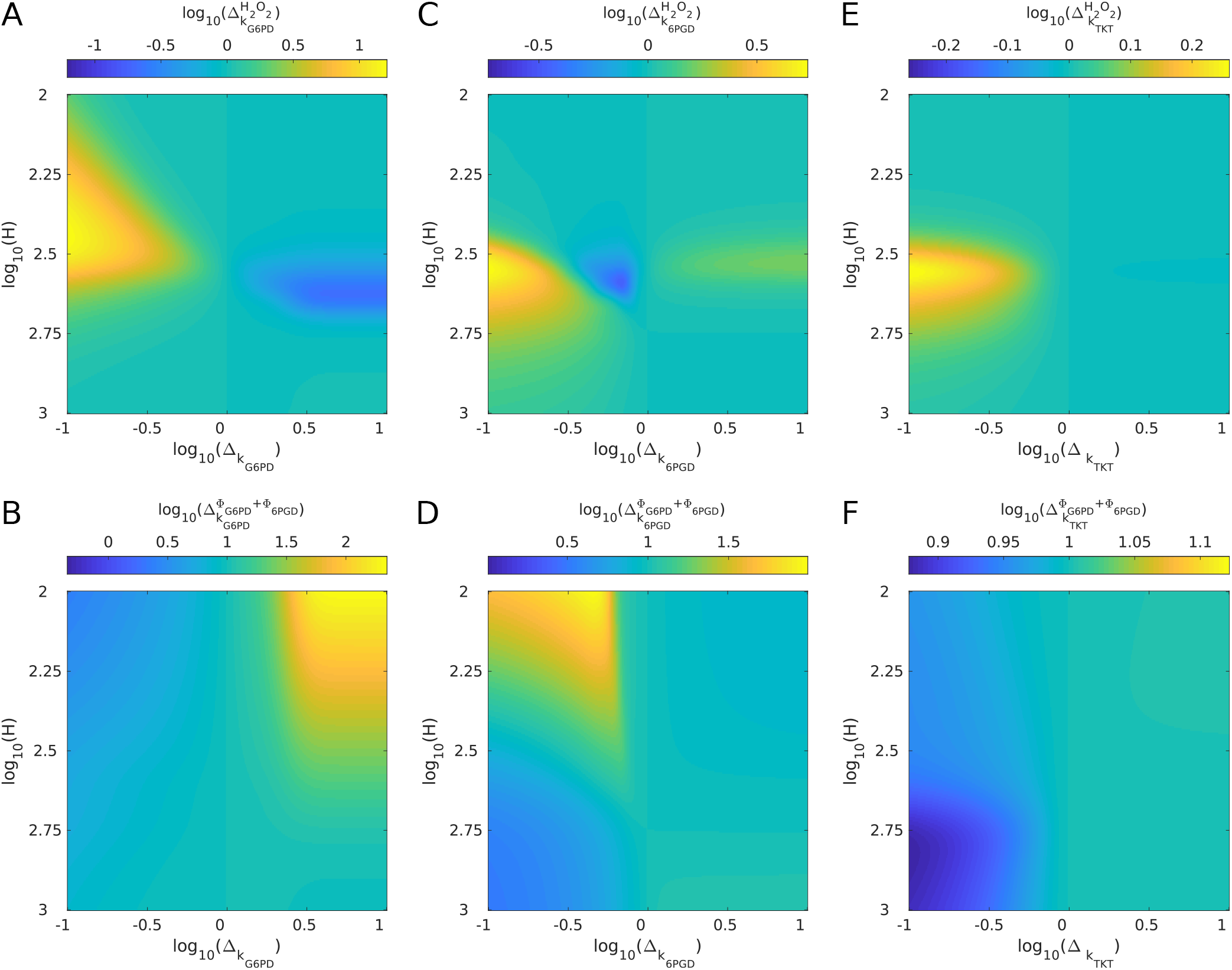
Gain/loss-of-function associated to modulated activity of PPP enzymes. Sensitivity 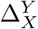 of output *Y* with respect to modulation of *X* (Eq 4) as a function of extracellular *H*_2_*O*_2_, H, and the extent of parameter modulation. The output variable *Y* is either *H*_2_*O*_2_ (A,C,E) and *NADPH*-producing flux (B,D,F) and the modulated parameter is either *k*_*G*6*P D*_ (A,B), *k*_6*P GD*_ (C,D) and *k*_*T KT*_ (E,F).

## Discussion

Inference and analysis of kinetic models of metabolism shed new light on the regulation of the oxidative stress response. Kinetic modeling is a natural mechanistic framework to identify or investigate metabolite-enzyme interactions (Link *et al*., 2013; Machado *et al*., 2015; Jahan *et al*., 2016; Reznik *et al*., 2017; Millard *et al*., 2017; Christodoulou *et al*., 2018). In such framework, regulation patterns have rather been investigated in large-scale manner through ensemble modeling allowing both parameters and regulatory structure to vary (Link *et al*., 2013; Christodoulou *et al*., 2018) or through metabolic control analysis (Millard *et al*., 2017). Instead, the present study focuses on a restricted subset of metabolite-enzyme interactions proposed to contribute to oxidative stress response, by performing careful setting, inference and perturbation analysis of regulatory kinetics. Such middle ground approach is well-suited to highlight cooperative mechanisms in complex regulation patterns. A selected set of allosteric regulatory mechanisms is not only required to explain experimental data, but is also demonstrated to exhibit a complementary range of efficiency and pleiotropic roles associated to the flux control of regulated reactions and concentration control of allosteric effectors. These modes of cooperation could be understand from a metabolic control perspective as second-order effects (Liebermeister, 2013), rather than those based on structural properties (Stelling *et al*., 2002; Notebaart *et al*., 2008; Sajitz-Hermstein and Nikoloski, 2013; Nikerel *et al*., 2012), optimality properties Wessely *et al*. (2011); Chubukov *et al*. (2012); Berkhout *et al*. (2012); Pfeuty and Thommen (2016) or dynamical properties (Krishna *et al*., 2007; Grimbs *et al*., 2007; Mulukutla *et al*., 2015) of metabolic networks.

In the context of oxidative stress, metabolic regulation is important to quickly reallocate metabolic fluxes to upregulate PPP-dependent production of *NADPH* as a cofactor of antioxidant enzymes (Mullarky and Cantley, 2015; Stincone *et al*., 2015; Dick and Ralser, 2015). Although data analysis and previous models have shown the involvement of several regulatory mechanisms in the metabolic response (Kuehne *et al*., 2017; Christodoulou *et al*., 2018), the present results refine our quantitative and mechanistic understanding of the contribution of regulation in the metabolic flux rerouting toward OxPPP. For low oxidative stress, the prominent regulatory mechanism involves the relief of *NADPH*-dependent inhibition of *G*6*PD* to boost PPP increase for a given stress-dependent consumption rate of *NADPH*. However, this mechanism is weakened by the concomittant decrease of G6P when OxPPP fluxes becomes of the order of downstream glycolytic flux. For higher oxidative stress, efficient upregulation of OxPPP fluxes must therefore coincide with an inhibition of glycolytic consumption of *G*6*P*, such as through the inhibition of *PGI*, to maintain *G*6*P* levels and enhance PPP response. For more severe oxidative stress, *H*_2_*O*_2_-dependent inhibition of *GAPD* is the only regulatory mechanism capable to reverse the glycolytic flux and allows for a cycling flux mode while the inhibition of PGI becomes ineffective. Although these three regulatory mechanisms contribute the most for different levels of intracellular *H*_2_*O*_2_, they nevertheless combine their effect sometimes with significant second-order synergy or redundancy effects. This is typically the case when for instance a given regulatory mechanism impacts the concentration response of the allosteric effectors of another regulatory mechanisms.

These three key modes of regulation also differ in the nature of allosteric regulators (*NADPH*, 6*PG* and *H*_2_*O*_2_) and feedback mechanisms. While *NADPH*-dependent and *H*_2_*O*_2_-dependent regulations implement a negative-feedback driven homeostasis mechanisms, 6*PG*-dependent inhibition of *PGI* generates a positive feedback mechanism which are much less common in metabolic pathways than feedback inhibition (Sauro, 2017; Alam *et al*., 2017; Locasale, 2018). Indeed, enhanced OxPPP flux leads to increased 6*PG* concentrations, such that stronger inhibition of *PGI* reduces *G*6*P* consumption and upregulates *G*6*PD* flux. Although this allosteric regulation has been reported in various studies (Gaitonde *et al*., 1989; Kuehne *et al*., 2015; Reznik *et al*., 2017), its functional role of this regulation has been seldomly debated (Vaseghi *et al*., 1999). Positive feedback induced by 6*PG*-dependent inhibition of *PGI* can lead to paradoxical behaviours in response to perturbation of 6*PGD* enzyme. The inhibition of 6*PGD* can (i) lower the maximal capacity flux and restrict flux usage for *NADPH* production, but can also (ii) further enhance 6*PG* increases and concomittant inhibition of glycolysis diverting flux into OxPPP. This ambivalent effect explains the contradicting experimental observations where genetic or pharmacological inhibition of 6*PGD* can either promote or decrease oxidative stress response depending on the context (Sun *et al*., 2019; Liu *et al*., 2019; Dubreuil *et al*., 2020). Another hypothesis to explore is that such positive feedback mediated by the upregulation of PPP intermediates can support discrete changes of metabolic phenotypes involving for instance the activation of AMPK also regulated by 6PG and R5P (Lin *et al*., 2015; Gao *et al*., 2019)

From a methodological perspective, this modeling study relies on a systematic procedure for model inference and parameter estimation, which deliberately considers more parameters than data to provide the possibility to update the probability distribution as additional data becomes available or as additional mechanistic hypothesis are tested. MCMC method is well adapted for such model updating procedure (Beck and Au, 2002; Lau and Gandy, 2014). Specifically, MCMC sampling can be restarted from a given parameter distribution with a modification of the score function and of the sample space. By testing the predictive capacity of the model on dataset that are not included in the model inference procedure, we could indeed identify sparse phenotypes poorly described by the model. Adding the corresponding dataset the monte-carlo sampling procedure is found to slightly shift the parameter set of solution to be consistent with these new dataset. Monte-Carlo methods can be combined with machine learning techniques to address metabolic control and optimality functionalities (Miskovic *et al*., 2019), providing an efficient framework for model selection. Such flexibility is valuable to extend our model to other model systems such as Erythrocytes (Thorburn and Kuchel, 1985) or microbes (Kerkhoven *et al*., 2013; Christodoulou *et al*., 2019), to hour-timescale response involving transcriptional regulation (Hayes and Dinkova-Kostova, 2014) or to other stress contexts such as glucose depletion and AMPK activation (Jeon *et al*., 2012; Lin *et al*., 2015).

## Materials and Methods

### Monte Carlo Markov Chain (MCMC) method

Monte Carlo Markov Chain (MCMC) method is used for estimating the distribution of (i) metabolic flux from ^13^C labeling data and (ii) kinetic model parameters from concentration data and flux estimation. These two computational problems are both defined by a set of *n*_*p*_ parameters 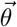, a set of *n*_*d*_ data values 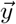 and a root mean square error 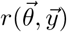 measuring the difference between data predicted by the parameters and the actual data (see Eqs 2 and 3). MCMC simulation methods use Monte Carlo sampling techniques to build Markov chains that converge to the posterior distribution of parameters *θ* associated to data *Y*. For both problems, we consider a bounded uniform distribution for the prior (see supplementary Table S2). For sampling, a Metropolis-Hasting algorithm is used to generate the random walk Markov chain based on (i) a gaussian jumping distribution and (ii) an acceptance rate function *α* that is the ratio of likelihood associated to the next parameter state and actual parameter state. Typically, the likelihood function can be written based on the expected or assumed distribution of observed data, as 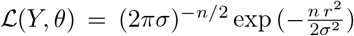. Specifically, we use the acceptance rate 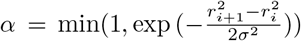 where the variances of jumping distribution and likelihood function *σ* are chosen to provide reasonable acceptance rate (> 10%) and convergence rate.

### SSA-based ^13^C-MFA

Stochastic simulation algorithm (SSA) for ^13^C-MFA is a direct method for the forward simulation problem to compute the dynamics and steady state of (mass) isotopomer distribution in isotopic labeling networks (Thommen *et al*., 2022). From a given flux distribution 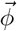 and initial labeling state, SSA computes the temporal evolution of isotopomer numbers which are pooled to obtain the mass isotopomer distribution *m*_*i,j*_ of species *j*. The sample size parameter of the algorithm is Ω = 100. From the experimental values of mass isotopomer 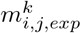 measured for *n*_*j*_ = 7 species *j* in *n*_*k*_ = 4 labeling conditions *k* (Kuehne *et al*., 2015), SSA is performed iteratively using a MCMC sampling method based on a random walk Metropolis algorithm for obtaining the posterior distribution of flux parameters. In such flux estimation procedure, the root mean square error function is given by:

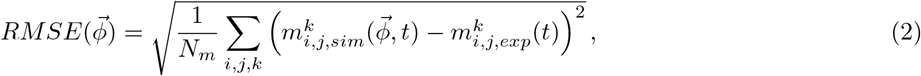

where the measurement time is *t* = 10mn and the number of experiments is *N*_*m*_ = *n*_*i*_*n*_*j*_*n*_*k*_ = 84.

### Kinetic model parameter estimation

The parameter estimation problem consists in scoring parameter set 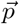 in the restricted parameter space 𝒫^*∗*^ using a normalized root mean square error (nRMSE) :

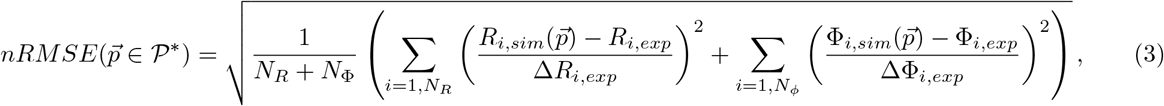

where 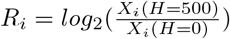 are concentration ratio in log scale and Φ_*i*_ are the estimated flux of reaction *i* in basal condition and in stress condition. Experimental data consists in *N*_*R*_ = 12 and *N*_Φ_ = 13 values of concentration ratio *R*_*i*_ and estimated flux Φ_*i*_ with a standard deviation Δ*R*_*i*_ = 1 and ΔΦ_*i*_ that is estimated from ^13^C-MFA.

This error function is used first with a population-based metaheuristic algorithm called evolution strategy to quickly explore and sample local solutions. Starting with a pool of *N*_*I*_ = 20 parameter vectors of random values (bounded uniform distribution), the algorithm involves three steps: (i) a reproduction step where a parent is randomly selected to be duplicated without applying any fitness criterion at this stage, and to generate *N*_*I*_ offspring; (ii) a mutation step where the parameters of each offspring are modified with a probability *p*_*m*_ through multiplication by a factor 10^*r*^, where *r* ∈ [−*a*_*m*_, *a*_*m*_] is a random number of uniform distribution; (iii) a selection step where the nRMSE of the *N*_*I*_ offspring are evaluated, and only the *N*_*I*_ highest-fitness individuals in the pool of 2*N*_*I*_ parameter sets (parents and offspring) are selected to generate the parents of the next generation. Finally, the optimization process terminates after a maximal number of generations (*N*_*G*_ = 10^4^) where a local minima of 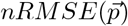 is usually reached. This evolution is repeated for a broad set of random initial conditions to obtain a first set of optimized models, some of which satisfies the plausibility criteria *nRMSE* < 1. The lowest-nRMSE model 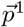 is used as initial condition of a MCMC algorithm used to sample a posterior probabilistic distribution of parameters. After a transient, a sample of 10^6^ accepted values are recorded to converge to a representative model ensemble associated to quasi-stationary distributions, while a random subsample of 10^5^ models is used for statistical analysis of model behaviors.

### Sensitivity analysis

Sensitivity quantities are defined to evaluate the effect of parameter changes on stress response behaviors. A model ensemble is defined by a set of model 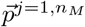. For a given kinetic model of parameter values 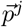, a parameter 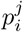 is multiplied by a factor Δ*p*_*i*_. A sensitivity factor that informs about the impact of a given parameter *p*_*i*_ on the output state variable *Y* in response to an extracellular level of hydrogen peroxide *H* is given by:

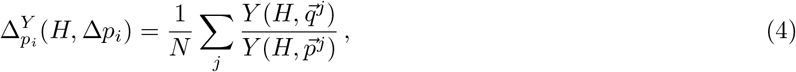

where 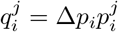 and *n*_*M*_ the number of model samples. The choice of parameter *p*_*i*_ and variation Δ*p*_*i*_ are selected to evaluate the impact of a regulatory mechanism (i.e., *p*_*i*_ are inhibitory parameters *Ki*_*i*_ and Δ*p*_*i*_ = ∞) or to mimmic overexpression or knockdown experiments (i.e., *p*_*i*_ are enzymatic activity rates *k*_*i*_ and 0.1 < Δ*p*_*i*_ < 10).

## Acknowledgments

This work has been supported by the LABEX CEMPI (ANR-11-LABX-0007) and by the Ministry of Higher Education and Research, Hauts de France council and European Regional Development Fund (ERDF) through the Contrat de Projets Etat-Region (CPER Photonics for Society P4S).

## Author Contributions

B.P. supervised the project; J.H., Q.T, F.A. B.P designed the methodology; J.H performed the simulations and analysis; J.H. and B.P. wrote the paper.

## Data and Code Availability

The Matlab code used in this study to perform model simulations and analysis are available at https://github.com/JHurb/HurbainPaper_Algorithm.

