## Supplementary Material for "Quantitative modeling of pentose phosphate pathway response to oxidative stress reveals a cooperative regulatory strategy"

*Abbreviations: G6P, glucose-6-phosphate; F6P, fructose-6-phosphate; FBP, fructose-1,6-bisphosphate; F2,6BP, fructose-2,6-bisphosphate; ALD, fructose 1,6 bisphosphate aldolase; DHAP, dihydroxyacetone phosphate; GAP, glyceraldehyde-3-phosphate; 6PG, 6-phosphogluconate; 6PGL, 6-phosphogluconolactone; X5P, xylulose 5-phosphate; R5P, ribose 5-phosphate; TPI, triosephosphate isomerase; Ru5P, ribulose 5-phosphate; OX, oxidative stress E4P, erythrose-4-phosphate; S7P, Sedoheptulose 7-phosphate; PPP, pentose phosphate pathway; HK, hexokinase; G6PD, G6P dehydrogenase; 6PGD, 6PG dehydrogenase; GLase, 6-phosphogluconolactonase; PRPP, Phosphoribosyl pyrophosphate; PGI, phosphoglucose isomerase; PFK(1), phosphofructokinase (type 1); FBPase, fructose-1,6-bisphosphatase; GAPD, GAP dehydrogenase; TLD, transaldolase; TKT, transketolase; ACC: Acetyl-CoA Carboxylase*

### 1 Differential equation Model of the metabolic network

#### 1.1 General formalism

The temporal behavior of the metabolic network is described by a differential equation system:

$$\frac{d\vec{x}}{dt} = \mathcal{S}\,\vec{\phi}(\vec{x}, H, \vec{p}) \tag{1}$$

- Vector components $x_i$ denote the concentration associated to metabolites $i$.
- $\mathcal{S}$ denotes the stochiometric matrix that represents a mapping of reaction rate vectors into a space of concentration time derivatives
- Vector components $\phi_j$ and $\phi_j(.)$ denote the rate and rate functions associated to the reaction catalyzed by the enzyme $j$. For reversible reactions, we also introduce directional fluxes satisfying the relation $\phi_j = \phi_j^+ - \phi_j^-$ where, by convention, the + direction goes from G6P to GAP in glycolysis and from R5P to GAP in nonoxidative PPP.
- Parameter vector $\vec{p}$ concatenates kinetic constants $k_i$, equilibrium constants $Keq_i$, Michaelis (saturation) constants $Km_i$ and inhibitory constants $Ki_i$, where $i$ denotes the enzyme.
- $\mathcal{S}\,\vec{\Phi}(\vec{X}, H, \vec{p}) = 0$ using capitalized letters for fluxes and concentrations corresponds to the steady-state equation.

#### 1.2 Metabolic network architecture

The set of enzymatic reactions and metabolites species considered in this study follows the common picture of the glycolytic, pentose phosphate, and gluthatione pathways, with a few noticeable simplifying assumptions:

- *Glucose metabolism:* Lower glycolysis, TCA cycle and other metabolic pathways are not considered with respect to available data restricted to upper glycolysis and pentose phosphate pathways.

- *Neglected cofactors:* Metabolite species do not include some organic or inorganic cofactors such as $ATP$, $Mg2+$ and $NAD+$ species, which are sometimes considered in kinetic models of glucose metabolism (Mulukutla *et al.*, 2015). We indeed assume that the variation of these metabolites is negligible during the early oxidative stress response or, if not, that their effect on the flux rerouting response is negligible compared to other mechanisms.

- *Neglected glycolytic reactions:* The reactions catalyzed by $PFK2$ and $FBPase2$ to convert between $F6P$ and $F2,6BP$ are not included as in many models (Vaseghi *et al.*, 1999; Mulukutla *et al.*, 2015), mainly because of the absence of data on the concentration of $F2,6BP$ metabolites which would lead to model parameter overfitting.

- *Merged NADP(H)-involved reactions:* Besides gluthatione pathways and PPP, the many other reactions involved $NADPH$ consumption and production (e.g., malic enzyme, IDH, fatty acid synthase, folate cycle) (Jeon *et al.*, 2012; Fan *et al.*, 2014; Gelman *et al.*, 2018; Chen *et al.*, 2019) are pooled in effective reactions called $NHN$ and $NNH$ respectively.

- *Neglected thioredoxin reactions:* The intracellular degradation of $H_2O_2$ is mediated by three main reaction systems namely catalase, gluthathione (via $GPx$) and thioredoxin (via $PRx$). In absence of data about redox state of $Prx$, we consider only gluthathione system, keeping in mind that $NADPH$ is the electron donor in both systems (Benfeitas *et al.*, 2014). The effect of $NADPH$-producing reaction on $H_2O_2$ are expected to be qualitatively similar for the two systems.

Accordingly, the set of equation reads:

| | | |
|---|---|---|
| $d[H_2O_2]/dt$ | $=$ | $\phi_{OX} - \phi_{CAT} - \phi_{GPx}$ |
| $d[GSH]/dt$ | $=$ | $2\phi_{GR} - 2\phi_{GPx}$ |
| $d[NADPH]/dt$ | $=$ | $\phi_{G6PD} + \phi_{6PGD} + \phi_{NNH} - \phi_{NHN} - \phi_{GR}$ |
| $d[G6P]/dt$ | $=$ | $\phi_{HK} - \phi_{G6PD} - \phi_{PGI}$ |
| $d[6PGL]/dt$ | $=$ | $\phi_{G6PD} - \phi_{6PGL}$ |
| $d[6PG]/dt$ | $=$ | $\phi_{6PGL} - \phi_{6PGD}$ |
| $d[Ru5P]/dt$ | $=$ | $\phi_{6PGD} - \phi_{RPI} - \phi_{RPE}$ |
| $d[X5P]/dt$ | $=$ | $\phi_{RPE} - \phi_{TKT1} - \phi_{TKT2}$ |
| $d[R5P]/dt$ | $=$ | $\phi_{RPI} - \phi_{TKT1} - \phi_{PRPP}$ |
| $d[S7P]/dt$ | $=$ | $\phi_{TKT1} - \phi_{TLD}$ |
| $d[E4P]/dt$ | $=$ | $\phi_{TLD} - \phi_{TKT2}$ |
| $d[GLC]/dt$ | $=$ | $\phi_{GLU} - \phi_{HK}$ |
| $d[F6P]/dt$ | $=$ | $\phi_{PGI} - \phi_{PFK} + \phi_{TAL} + \phi_{TKT2}$ |
| $d[FBP]/dt$ | $=$ | $\phi_{PFK} - \phi_{ALD}$ |
| $d[DHAP]/dt$ | $=$ | $\phi_{ALD} - \phi_{TPI}$ |
| $d[GAP]/dt$ | $=$ | $\phi_{ALD} + \phi_{TKT1} - \phi_{TAL} + \phi_{TKT2} + \phi_{TPI} - \phi_{GAPD}$ |

**Table S1: Equation table.**

#### 1.3 Regulatory architecture

We focus on the set of regulation that are expected or acknowledged to contribute to the early ($< 10mn$) metabolic response based on biochemical and functional evidences in Mammalian cells:

- $NADPH$-dependent inhibition of $G6PD$ and $6PGD$ (Yoshida and Lin, 1973; Holten *et al.*, 1976; Christodoulou *et al.*, 2018).

- $H_2O_2$-dependent inhibition of $GAPD$ (Peralta *et al.*, 2015; van der Reest *et al.*, 2018).
- $6PG$-dependent inhibition of $PGI$ (Parr, 1956; Kahana *et al.*, 1960; Gaitonde *et al.*, 1989; Kuehne *et al.*, 2015; Dubreuil *et al.*, 2020).
- Inhibition of $NADPH$ consumption through rapid shutdown of biosynthetic pathways involving for instance $ACC$ (Jeon *et al.*, 2012; Fan *et al.*, 2014).

In contrast, we neglect genome-scale regulation occurring at slower time scales ($> 10mn$) and posttranslation regulation activated by glucose depletion (e.g., via $AMPK$). For comparison, previous models of oxidative stress response does not consider any regulation (Kerkhoven et al, 2013) or does consider all possible metabolite-enzyme interactions (Christodoulou *et al.*, 2018). This intermediate strategy of restricting to a few set of regulation is driven both by the constraint of parameter identifiability (avoid overfitting) and the goal of understanding an well-supported regulatory pattern rather than exploring or discovering regulatory patterns.

##### 1.4 Reaction kinetics scheme

Reaction rates can be described by diverse kinetic laws (e.g., mass action, Michaelis-Menten, Hill, Monod-Wyman-Changeux...). In the present work, we mainly use the generalized mass action kinetics, eventually associated with inhibition:

$$MA(m,n): \ \phi(S,P,I,k_i,Keq_i,Ki_i) = k_i \left( \prod_{j=1,m} S_j - (Keq_i)^{-1} \prod_{j=1,n} P_j \right) (1 + I/Ki_i)^{-1} \tag{2}$$

where $m$ and $n$ are the number of substrates and products. However in the context of oxidative stress response, the significant increase of some metabolite concentrations may require to take into account the impact of saturated enzymatic activity using Michaelis-Menten-like kinetics. The generalized irreversible and reversible uni/bi-substrate reaction kinetics (Rohwer *et al.*, 2006):

$$MM(m,0) \quad : \quad \phi(S,P,k_i,Km_i) = \frac{k_i \prod_j S_j}{\prod_j (1 + S_j/Km_{sj,i})} \tag{3}$$

$$MM(m,n) \quad : \quad \phi(S,P,k_i,Keq_i,Km_i) = \frac{k_i \left( \prod_j S_j - (Keq_i)^{-1} \prod_j P_j \right)}{\prod_j (1 + S_j/Km_{sj,i} + P_j/Km_{pj,i})} \tag{4}$$

where $j = 1$ for unisubstrate reactions and $j = 1,2$ for bisubstrate reactions. In some bisubstrate reactions, the assumption that $S \ll K_m$ is made so as to reducing the number of terms and parameter in denominator. Specifically, Michaelis constants are considered only for $6PG$, $H_2O_2$, $GSSG$ and $GSH$ and in nonoxidative PPP reactions. For $GPx$ and $GR$ reactions, the saturation terms follow a common scheme based on experimental data (Benfeitas *et al.*, 2014). A minimal description of saturation in nonoxidative PPP reactions consists in keeping only product terms (which prevails in case where $S_i/Km_i$ larger than one) and assume a same Michaelis constant for substrates and products.

| Reaction | Law | Rate equation |
|---|---|---|
| $\phi_{OX}$ | T | $k_{OX} + k_{diff}([H_2O_2]_{ext} - [H_2O_2])$ |
| $\phi_{CAT}$ | MA(1,0) | $k_{CAT}[H_2O_2]$ |
| $\phi_{GPx}$ | MM(2,0) | $k_{GPx}[H_2O_2][GSH]/(\frac{[GSH]}{KmG_{GPx}} + \frac{[H2O2]}{KmH_{GPx}})$ |
| $\phi_{GR}$ | MM(2,0) | $k_{GR}[NADPH][GSSG]/(1 + \frac{[GSSG]}{KmG_{GR}} + \frac{[NADPH]}{KmN_{GR}})$ |
| $\phi_{NNH}$ | MA(1,0) | $k_{NAD}[NADP^{+}]$ |
| $\phi_{NHN}$ | MA(1,0)+CI | $k_{NADPH}[NADPH]/(1 + \frac{[H_2O_2]}{Ki_{NNH}})$ |
| $\phi_{G6PD}$ | MA(2,0)+CI | $k_{G6PD}[G6P][NADP^{+}]/(1 + \frac{[NADPH]}{Ki_{G6PD}})$ |
| $\phi_{6PGL}$ | MA(1,0) | $k_{GLase}[6PGL]$ |
| $\phi_{6PGD}$ | MM(2,0)+CI | $k_{6PGD}[6PG][NADP^{+}]/(1 + \frac{[NADPH]}{Ki_{6PGD}} + \frac{[6PG]}{Km_{6PGD}})$ |
| $\phi_{RPI}$ | MA(1,1) | $k_{RPE}([Ru5P] - \frac{[X5P]}{Keq_{RPE}})$ |
| $\phi_{RPE}$ | MA(1,1) | $k_{RPI}([Ru5P] - \frac{[R5P]}{Keq_{RPI}})$ |
| $\phi_{PRPP}$ | MM(1,0) | $k_{PRPP}[R5P]/(1 + \frac{[R5P]}{Km_{PRPP}})$ |
| $\phi_{TKT1}$ | MM(2,2) | $k_{TKT1}([R5P][X5P] - \frac{[GAP][S7P]}{Keq_{TKT1}})/(1 + \frac{([R5P][X5P]}{Km_{TKT1}} + \frac{[GAP][S7P])}{Km_{TKT1}})$ |
| $\phi_{TLD}$ | MM(2,2) | $k_{TLD}([GAP][S7P] - \frac{[F6P][E4P]}{Keq_{TLD}})/(1 + \frac{([GAP][S7P]}{Km_{TLD}} + \frac{[F6P][E4P])}{Km_{TLD}})$ |
| $\phi_{TKT2}$ | MM(2,2) | $k_{TKT2}([E4P][X5P] - \frac{[F6P][GAP]}{Keq_{TKT2}})/(1 + \frac{([E4P][X5P]}{Km_{TKT2}} + \frac{[F6P][GAP]}{Km_{TKT2}})$ |
| $\phi_{HK}$ | MA(1,0)+CI | $k_{HK}[GLC]/(1 + \frac{[G6P]}{Ki_{HK}})$ |
| $\phi_{PGI}$ | MA(1,1)+CI | $k_{PGI}([G6P] - \frac{[F6P]}{Keq_{PGI}})/(1 + \frac{[6PG]}{Ki_{PGI}})$ |
| $\phi_{PFK}$ | MA(1,0) | $k_{PFK}[F6P] - k_{FBPase}[FBP]$ |
| $\phi_{ALD}$ | MA(1,2) | $k_{ALD}([FBP] - \frac{[DHAP][GAP]}{Keq_{ALD}})$ |
| $\phi_{TPI}$ | MA(1,1) | $k_{TPI}([DHAP] - \frac{[GAP]}{Keq_{TPI}})$ |
| $\phi_{GAPD}$ | MA(1,0)+CI | $k_{GAPD}[GAP]/(1 + \frac{[H_2O_2]}{Ki_{GAPD}})$ |

**Table S2: Reaction table.** CI: competitive inhibition. MM: Michaelis-Menten with some order approximation (*); MA: Mass-action;

| Parameter | Value | Range | References |
|---|---|---|---|
| $k_{GPx}$ | 1s$^{-1}$ | $[10^{-1}; 10^{1}]$ | (Benfeitas *et al.*, 2014) |
| $KmH_{GPx}$ | 0.04$\mu$M | $[10^{-1}; 10^{1}]$ | (Benfeitas *et al.*, 2014) |
| $KmG_{GPx}$ | 9.72$\mu$M | $[10^{-1}; 10^{1}]$ | (Benfeitas *et al.*, 2014) |
| $k_{GR}$ | 49s$^{-1}$ | $[10^{-1}; 10^{1}]$ | (Benfeitas *et al.*, 2014) |
| $KmN_{GR}$ | 8.5$\mu$M | $[10^{-1}; 10^{1}]$ | (Benfeitas *et al.*, 2014) |
| $KmG_{GR}$ | 65$\mu$M | $[10^{-1}; 10^{1}]$ | (Benfeitas *et al.*, 2014) |
| $Ki_{G6PD}$ | 10$\mu$M | $[10^{-1}; 10^{1}]$ | (Yoshida and Lin, 1973; Kuehne *et al.*, 2015) |
| $Ki_{6PGD}$ | 10$\mu$M | $[10^{-1}; 10^{1}]$ | (Yoshida and Lin, 1973; Holten *et al.*, 1976) |
| $Ki_{PGI}$ | 100$\mu$M | $[10^{-1}; 10^{1}]$ | (Kuehne *et al.*, 2015) |
| $Ki_{GAPD}$ | 100$\mu$M | $[10^{-1}; 10^{1}]$ | (Peralta *et al.*, 2015) |
| $Ki_{NNH}$ | 100$\mu$M | $[10^{-1}; 10^{1}]$ | |
| $Km_{6PGD}$ | 50$\mu$M | | (Ceyhan *et al.*, 2005; Liu *et al.*, 2019) |
| $Km_{PRPP}$ | 65$\mu$M | | (Hove-Jensen *et al.*, 2017) |
| $Keq_{RPE}$ | 1.68 | | (Li *et al.*, 2011) |
| $Keq_{RPI}$ | 1.23 | | (Li *et al.*, 2011) |
| $Keq_{TKT1}$ | 1.62 | | (Li *et al.*, 2011) |
| $Keq_{TLD}$ | 0.36 | | (Li *et al.*, 2011) |
| $Keq_{TKT2}$ | 30 | | (Li *et al.*, 2011) |
| $Keq_{PGI}$ | 0.34 | | (Li *et al.*, 2011) |
| $Keq_{ALD}$ | 66$\mu$M | | (Li *et al.*, 2011) |
| $Keq_{TPI}$ | 19.2 | | (Li *et al.*, 2011) |
| $k_{diff}$ | 1s$^{-1}$ | $[10^{-1}; 10^{1}]$ | (Benfeitas *et al.*, 2014) |
| $\Phi_{glu}$ | 40$\mu$M.s$^{-1}$ | | |
| $[NADP(H)]_{tot}$ | 30$\mu$M | | (Benfeitas *et al.*, 2014) |
| $[GSH]_{tot}$ | 3mM | | (Benfeitas *et al.*, 2014) |
| $k_i$ | 1s$^{-1}$ | $[10^{-4}; 10^{1}]$ | |

**Table S3: Parameter space and values.** Range defines the range of variation around the indicated value. Blank range indicates fixed value. Some values (e.g., $Km$ or $Ki$, not $Keq$) are approximated to rounded or averaged values from experimental measurements

.


Gelman, S. J. *et al.* (2018). Consumption of NADPH for 2-HG synthesis increases pentose phosphate pathway flux and sensitizes cells to oxidative stress. *Cell Rep.*, **22**(2), 512–522.

Holten, D. *et al.* (1976). Regulation of pentose phosphate pathway dehydrogenases by NADP+/NADPH ratios. *Biochem. Biophys. Res. Commun.*, **68**(2), 436–441.

Hove-Jensen, B. *et al.* (2017). Phosphoribosyl diphosphate (PRPP): biosynthesis, enzymology, utilization, and metabolic significance. *Microbiol. Mol. Biol. Rev.*, **81**(1), e00040–16.

Jeon, S.-M. *et al.* (2012). AMPK regulates NADPH homeostasis to promote tumour cell survival during energy stress. *Nature*, **485**(7400), 661–665.

Kahana, S. E. *et al.* (1960). The kinetics of phosphoglucoisomerase. *J. Biol. Chem.*, **235**(8), 2178–2184.

Kuehne, A. *et al.* (2015). Acute Activation of Oxidative Pentose Phosphate Pathway as First-Line Response to Oxidative Stress in Human Skin Cells. *Mol. Cell*, **59**(3), 359–371.

Li, X. *et al.* (2011). A database of thermodynamic properties of the reactions of glycolysis, the tricarboxylic acid cycle, and the pentose phosphate pathway. *Database*, **2011**, 1–15.

Liu, R. *et al.* (2019). Tyrosine phosphorylation activates 6-phosphogluconate dehydrogenase and promotes tumor growth and radiation resistance. *Nat. Commun.*, **10**(1), 1–14.

Mulukutla, B. C. *et al.* (2015). Multiplicity of steady states in glycolysis and shift of metabolic state in cultured mammalian cells. *PLoS One*, **10**(3), e0121561.

Parr, C. (1956). Inhibition of phosphoglucose isomerase. *Nature*, **178**(4547), 1401–1401.

Peralta, D. *et al.* (2015). A proton relay enhances H2O2 sensitivity of GAPDH to facilitate metabolic adaptation. *Nat. Chem. Biol.*, **11**(2), 156–163.

Rohwer, J. *et al.* (2006). Evaluation of a simplified generic bi-substrate rate equation for computational systems biology. *IET Syst Biol*, **153**(5), 338–341.

van der Reest, J. *et al.* (2018). Proteome-wide analysis of cysteine oxidation reveals metabolic sensitivity to redox stress. *Nat. Commun.*, **9**(1), 1–16.

Vaseghi, S. *et al.* (1999). In vivo dynamics of the Pentose Phosphate Pathway in Saccharomyces cerevisiae. *Metab. Eng.*, **1**(2), 128–140.

Yoshida, A. and Lin, M. (1973). Regulation of glucose-6-phosphate dehydrogenase activity in red blood cells from hemolytic and nonhemolytic variant subjects. *Blood*, **41**(6), 877–891.

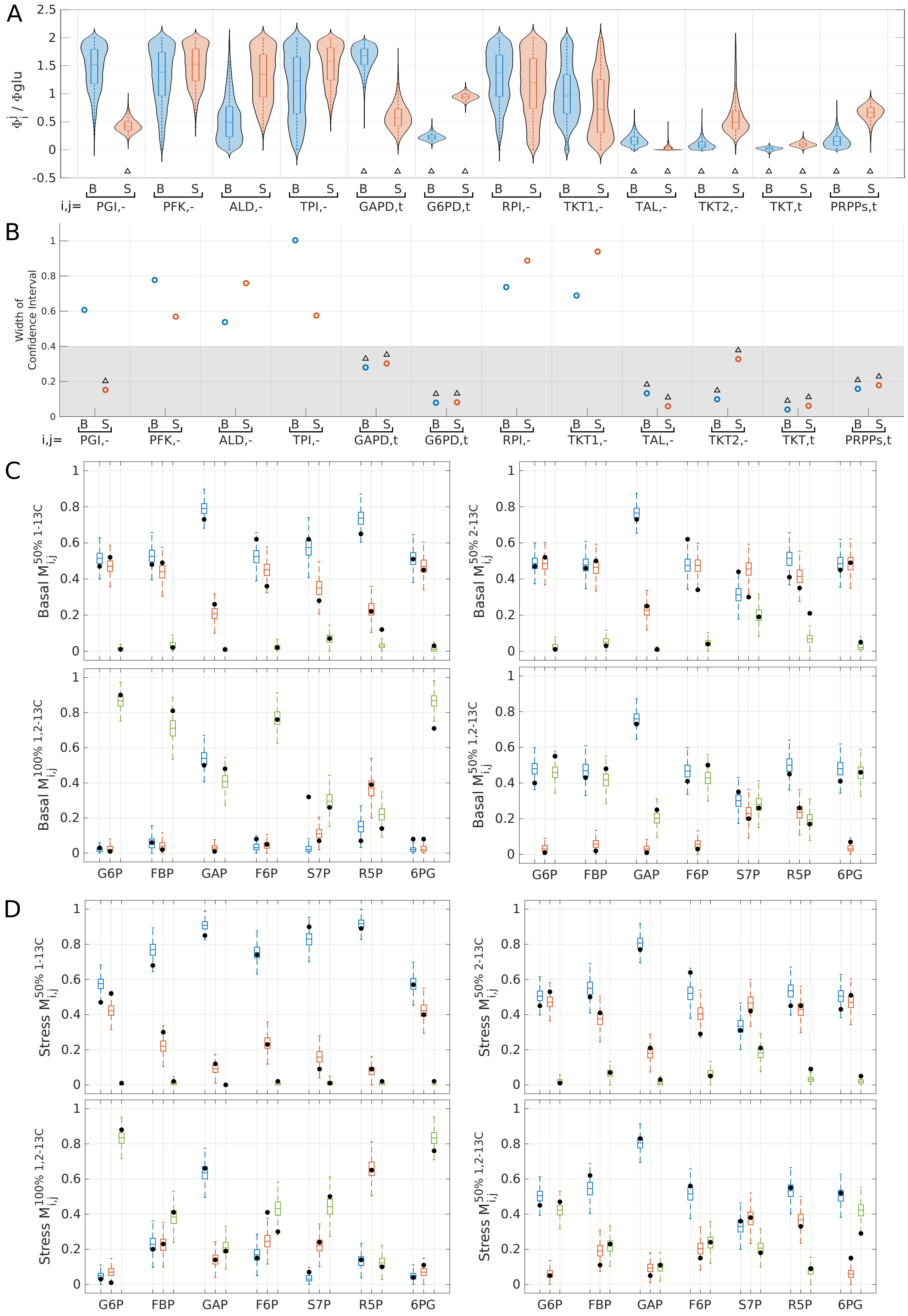


**Figure S1: Estimated flux distribution obtained with MCMC sampling.** (A) Violin plot of the posterior flux distribution obtained from MCMC sampling of SSA simulation of the labeled isotopic systems. Blue and red indicates basal and stress conditions. (B) 50% Confidence intervals associated with these distribution determines the specific set of flux (indicated by triangles) for which estimation is accurate and narrow enough to be used for kinetic model fitting. (C,D) Box plot of $M_{i,j}$ obtained from MCMC sampling of SSA simulation of the labeled isotopic systems in four labeling conditions as compared with experimental values (black circles) in absence (C) and presence (D) of oxidative stress respectively.

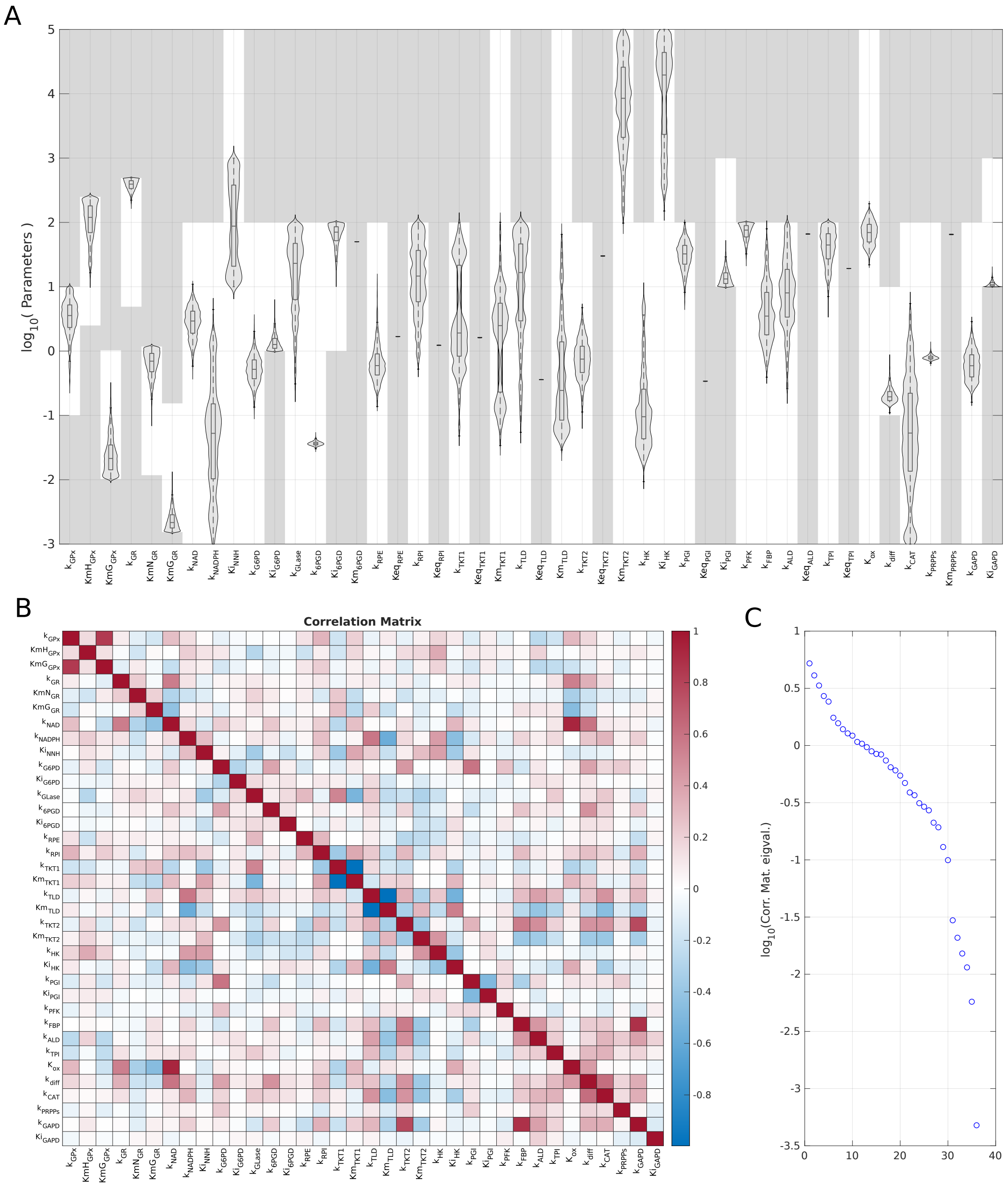


**Figure S2: Analysis of parameter distributions.** (A) Violin plot of parameter distribution obtained through MCMC sampling. White domains indicates the uniform prior distribution of parameters used for MCMC sampling. (B) Correlation matrix excluding fixed value parameters. (C) Ranked eigenvalues of the correlation matrix. The 5 highest eigenvalues contributes to 50% of the variance (ie, 50% of the variance can be captured by four linear combinations of parameters). Eigenvectors involving a small number of parameters are not observed which restrains the possibility for further parameter number reduction.

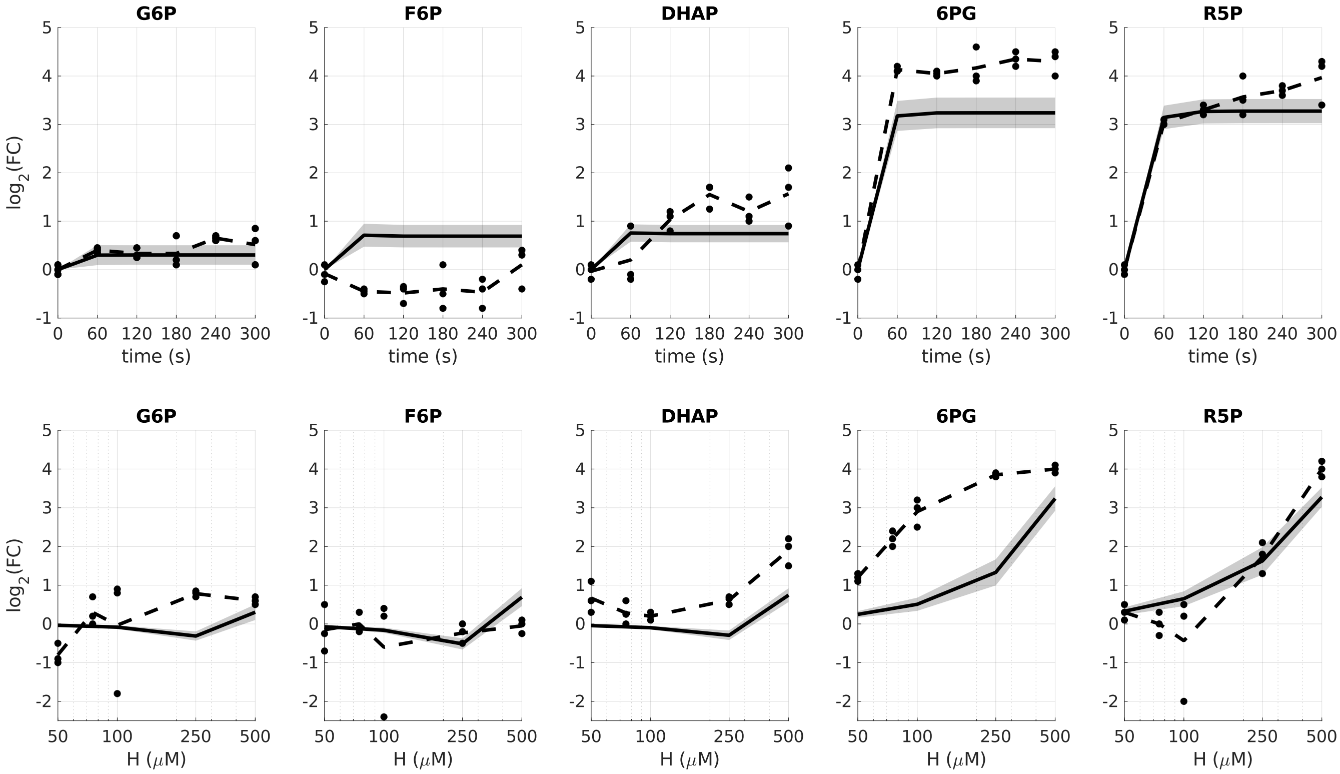


**Figure S3: Model prediction test.** Dynamical and dose-reponse behaviors of the model ensemble $\mathcal{P}^{opt}$ as compared with experimental data in (Kuehne *et al.*, 2015) that were not used as target dataset in the adjustement score function for model parameter estimation. FC means fold-change of metabolites concentration after stress with respect values to just before as function of measurement time (up panels) and peroxide concentration.

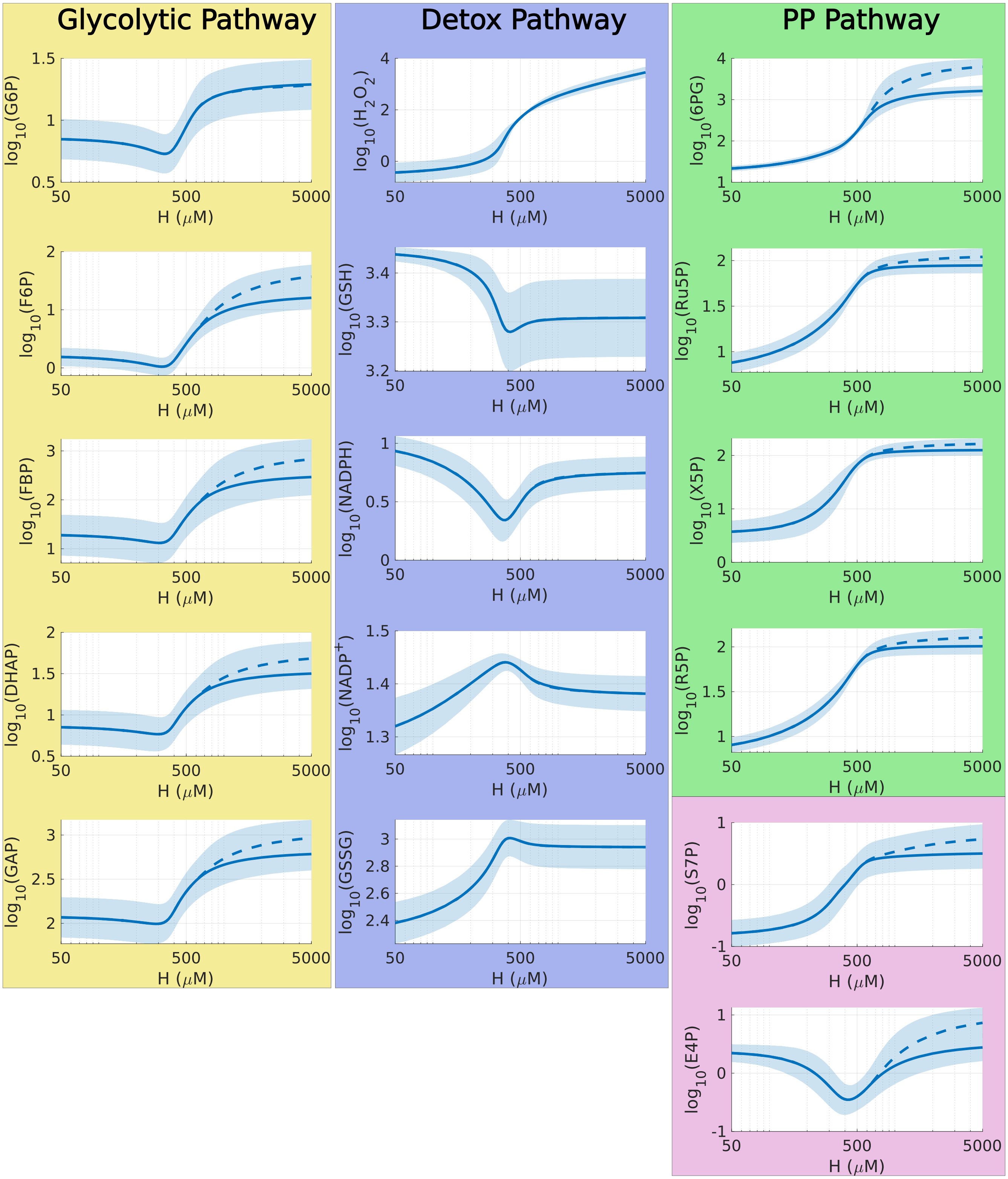


**Figure S4: Dose response of metabolite concentrations.** Complementary data associated to Figure 5B including all metabolite species.

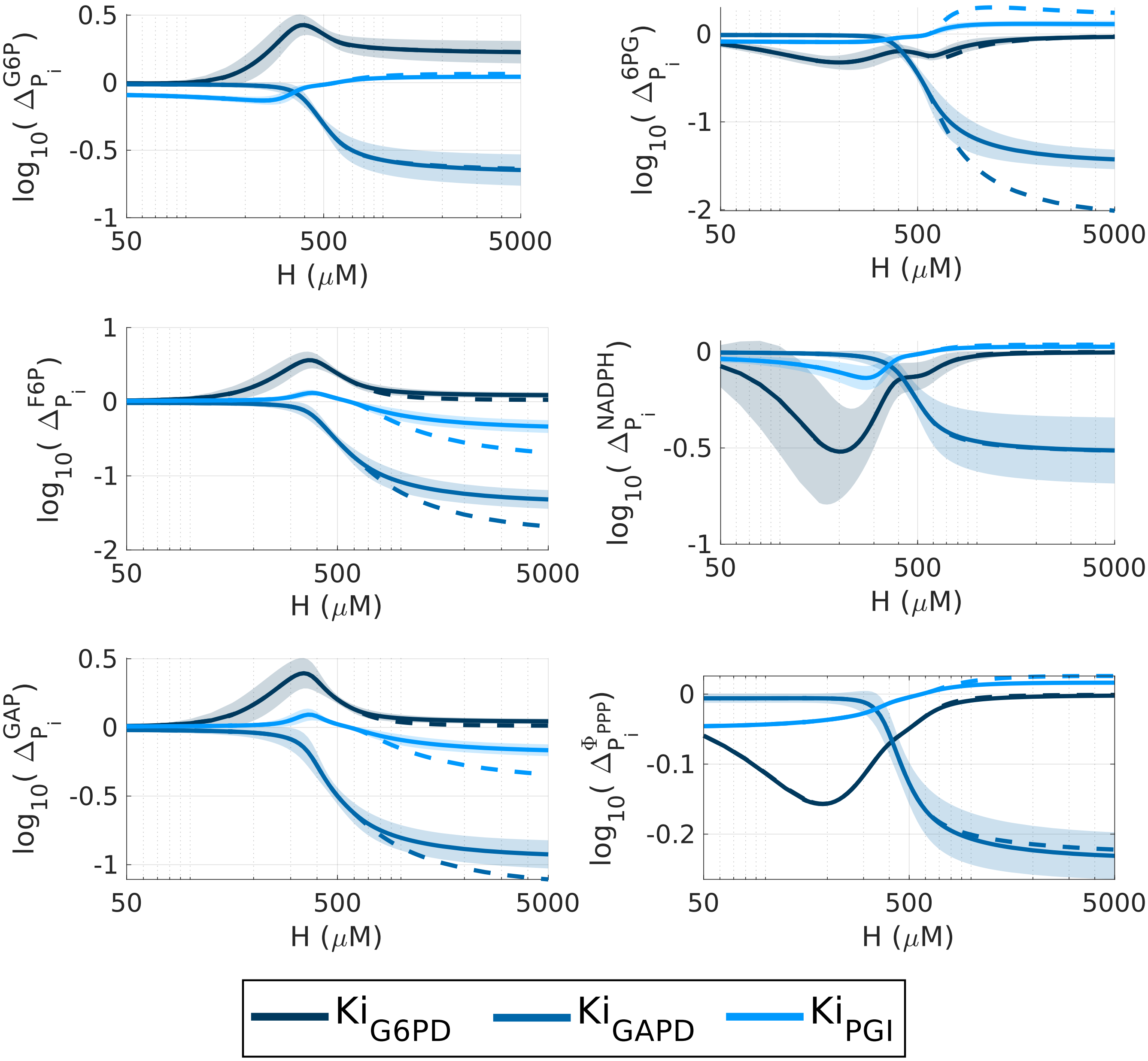


**Figure S5: Regulation analysis based on gain/loss-of-function simulations.** Complementary data associated to Figure 6. An opposite responses of glycolytic metabolites G6P, F6P and GAP (positive versus negative) is observed between the deletions of PPP inhibition ($Ki_{G6PD}$) and of glycolytic inhibition ($Ki_{PGI}$,$Ki_{GAPD}$), which explains some second-order effects.

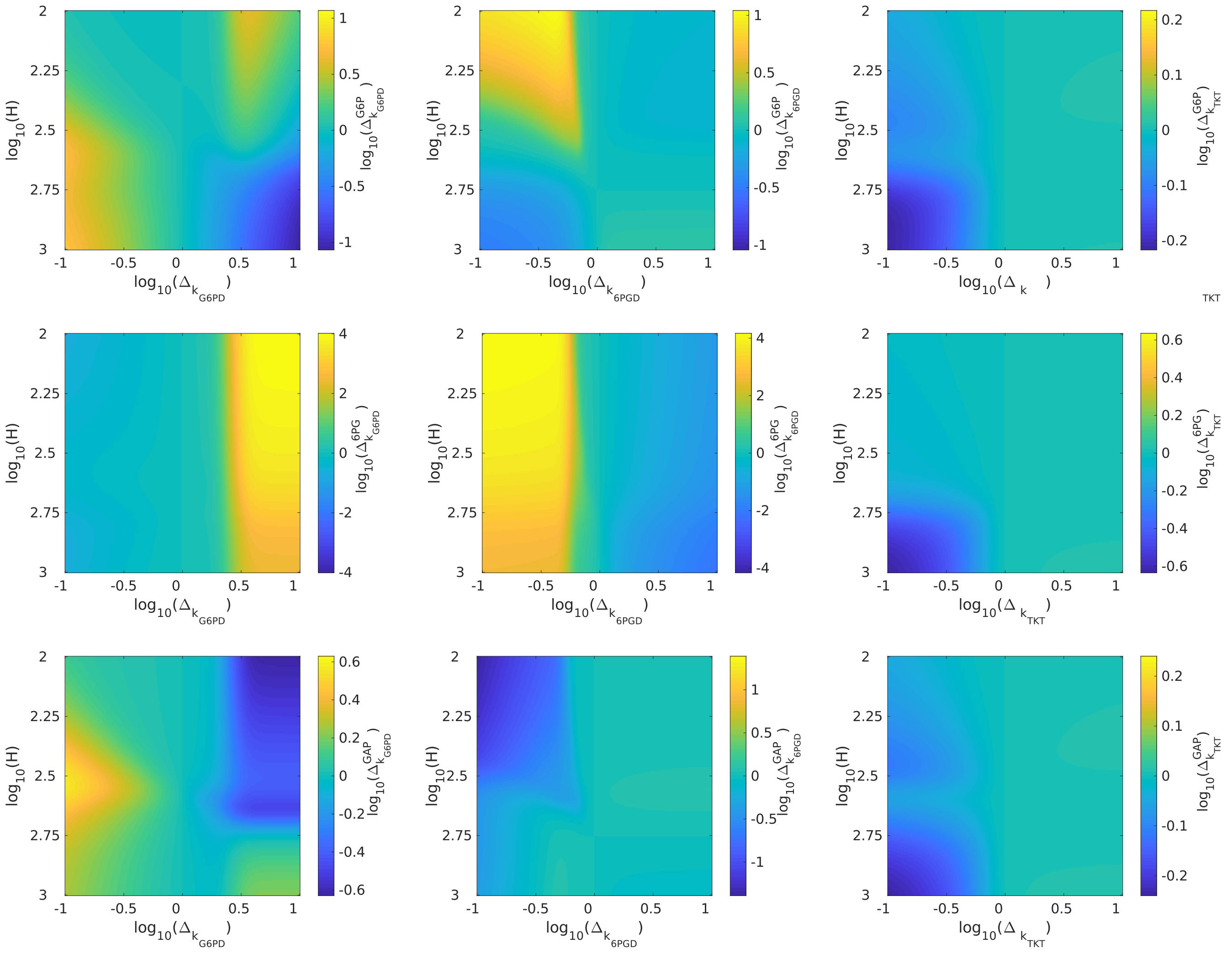


**Figure S6: Changes of key metabolite concentrations (G6P, 6PG and GAP) associated to modulated activity of PPP enzymes and oxidative stress level.** Complementary data associated to Figure 7. These can be seen as non-infinitesimal and non-normalized concentration control coefficients. Downregulation of $k_{G6PD}$ shows increased G6P and GAP and decreased 6PG (consistently with Fig.3 of (Kuehne *et al.*, 2015).) Downregulation of $k_{TKT}$ shows decreased G6P, GAP and G6P (consistently with Fig.3 of (Kuehne *et al.*, 2015) except for 6PG) Downregulation of $k_{6PGD}$ shows an opposite qualitative response to that of $k_{G6PD}$.
